# The genomic consequences and persistence of sociality in spiders

**DOI:** 10.1101/2024.04.22.590577

**Authors:** Jilong Ma, Jesper Bechsgaard, Anne Aagaard, Palle Villesen, Trine Bilde, Mikkel Heide Schierup

## Abstract

In cooperatively breeding social animals, a few individuals account for all reproduction. In some taxa, sociality is accompanied by a transition from outcrossing to inbreeding, in concert, these traits act to reduce effective population size, potentially rendering transitions to sociality ‘evolutionarily dead-ends’. We addressed this hypothesis in a comparative genomic study in spiders, where sociality has evolved independently at least 23 times, but social species are recent and evolutionarily short-lived. We present genomic evidence for the evolutionary dead-end hypothesis in three independent transitions to sociality in the spider genus *Stegodyphus*. We sequenced, assembled and annotated high-quality, chromosome-level reference genomes from three pairs of closely related social and subsocial *Stegodyphus* species. Genome sizes range from 2.65 Gb to 3.32 Gb with high synteny, and we identify 10,065 single-copy orthologous genes. We timed the divergence between the social and subsocial species pairs to be from 1.3 to 1.8 million years. Social evolution in spiders involves a shift from outcrossing to inbreeding and from equal to female-biased sex ratio, causing severe reductions in effective population size and decreased efficacy of selection. Based on analyses of purifying selection, we determined whether transitions to sociality co-occurred with divergence. We show that transitions to sociality only had full effect on purifying selection at 119 kya (95CI: 71 kya -169 kya), 260 kya (95CI: 231 ky - 289 kya) and 279 kya (95CI: 230 kya - 332 kya) respectively, and follow remarkably similar convergent trajectories of progressive loss of diversity and shifts to an increasingly female-biased sex ratio. This almost deterministic genomic response to sociality may explain why social spider species do not persist. What causes species extinction is not clear, but could be either selfish meiotic drive that eliminates the production of males, or an inability to retain genome integrity in the face of extremely reduced efficacy of selection.

## Introduction

The evolution of cooperative breeding, where some individuals forego their own reproduction to become helpers that assist other individuals in raising the offspring, is widespread in the animal kingdom (Rubenstein and Abbot 2017; Alexander 1974). In the most extreme cases, as in eusocial insects, strict division of labour results in highly complex societies with one or a few reproducing queens and countless non-reproducing helpers that are further organised into specialised sterile worker castes that each have specific tasks to perform(Crespi and Yanega 1995). In most other cooperatively breeding taxa, individuals can switch between roles of reproducer and helper depending on age, hierarchy and ecological conditions (Alexander 1974; Koenig et al. 1992; Ross and Keller 1995; Faulkes et al. 1997; Lubin and Bilde 2007). Cooperative breeding has evolved independently in divergent taxa including insects, arachnids, birds, mammals and fish. Common to all social cooperative species is that the evolution of cooperative breeding occurs in family groups and that helpers gain inclusive fitness benefits by assisting related individuals in reproducing, highlighting the importance of relatedness and kin selection in the evolution of sociality(Hamilton 1964).

As only a subset of individuals accounts for all reproduction in the social group, cooperative breeding invariably implies a reduction in effective population size (Settepani et al. 2017; Romiguier et al. 2014; Nomura and Takahashi 2012). Most cooperatively breeding species maintain a predominantly outcrossing mating system governed by pre-mating or sex-biased dispersal that takes place prior to mating. However, in a subset of species, the transition to sociality is accompanied by a switch from outcrossing to strict inbreeding through the elimination of pre-mating dispersal and reproduction among highly related group members. These taxa show a ‘social syndrome’ characterised by convergent independent transitions to cooperative breeding, regular inbreeding, reproductive skew (i.e. a subset of adult females reproduce), and a primary female biased sex ratio(Bilde and Lubin 2011; Settepani et al. 2017; Vanthournout et al. 2018). This combination of traits has evolved convergently multiple times in spiders (Lubin and Bilde 2007), thrips (Chapman et al. 2000), aphids (Johnson et al. 2000), beetles (Keller et al. 2011) and possibly also mole rats(Reeve et al. 1990). In concert, these traits further act to reduce effective population sizes, and this is predicted to have negative consequences for population genetic diversity and the efficacy of selection through elevated genetic drift (Charlesworth and Wright 2001; Charlesworth 2003; Bechsgaard et al. 2019).

Spiders are an evolutionarily old group (>300 million years) (Garwood et al. 2016), and signs of social behavior were reported from fossil evidence in 100 million-year-old amber (Poinar and Buckley 2012). Although sociality in spiders is rare, the combination of cooperative breeding and inbreeding mating system has evolved independently at least 23 times (Lubin and Bilde 2007) in 6 spider families, constituting ∼23 of more than 50,000 extant spider species (World Spider Catalog). Phylogenetic analyses show that social spider lineages are evolutionarily young as they are found at the terminal branches. Furthermore, there are no described cases of speciation following transitions to sociality (Lubin and Bilde 2007; Agnarsson et al. 2006), in contrast to the social insects, where relatively few transitions to sociality are followed by rich diversifications (Danforth 2002; Moreau et al. 2006). These phylogenetic patterns substantiate the hypothesis that the joint transitions to sociality and inbreeding may represent an “evolutionary dead-end”(Takebayashi and Morrell 2001; Settepani et al. 2017), where social transitions are driven by ecological benefits, while long term costs of repeated inbreeding cause the demise of social lineages.

Sociality in spiders involves living in communal nests where they cooperate in web building, prey capture, and offspring care (Avilés 2021). Although there is a degree of behavioral differentiation (Settepani et al. 2013), they do not show eusociality like in ants and bees (Crespi and Yanega 1995), as there are no casts or strict division of labor (Lubin and Bilde 2007). Phylogenetic relationships indicate that sociality in spiders evolves from subsocial ancestors (Kullmann 1972; Avilés 2010; Agnarsson et al. 2006). In subsocial species, females provide extensive maternal care for their offspring over a prolonged period after hatching (Lubin and Bilde 2007; Grinsted et al. 2014), and the juveniles of subsocial spider species spend the first weeks in the natal nest before they disperse to establish a solitary life (Bilde et al. 2005). While both social and subsocial species show elaborate maternal care of the offspring, subsocial species maintain premating dispersal and consequently an outcrossing mating system (Bilde et al. 2005). In contrast, sociality is associated with the complete elimination of premating dispersal, which results in obligatory inbreeding as siblings reproduce within the group (Lubin and Bilde 2007). This is accompanied by the evolution of female biassed sex ratios (Vanthournout et al. 2018; Bechsgaard et al. 2019), cooperative breeding where female helpers provide extensive allomaternal care (Junghanns et al. 2017, 2019) and reproductive skew (Lubin and Bilde 2007). The convergent evolution of traits that constitute the ‘social syndrome’ and the phylogenetic relationships between social and subsocial species makes this system ideal for understanding causes and consequences of transitions to sociality and inbreeding.

In the spider genus *Stegodyphus*, which contains more than 25 species, sociality has evolved three times independently (Johannesen et al. 2007; Settepani et al. 2016), and each social species has a closely related subsocial sister-species (see Fig 1A) suggesting that sociality evolved recently (Bechsgaard et al. 2019; Settepani et al. 2016). The social *Stegodyphus* species express the “social syndrome” characteristics, which are predicted to reduce their effective population sizes dramatically (Bechsgaard et al. 2019) which increases the strength of drift and reduces the efficacy of natural selection (Kimura et al. 1963). This substantiates the question of whether consequences of low Ne impacts on the evolutionary persistence of social lineages. Reports of substantially reduced neutral genetic diversity (Settepani et al. 2017) and elevated rates of nonsynonymous to synonymous substitution rates (Mattila et al. 2012; Settepani et al. 2016; Bechsgaard et al. 2019, 2022) in social *Stegodyphus* species are consistent with reduced effective population sizes. Repeated inbreeding over evolutionary time is likely to have resulted in purging but also elevated rates of fixation of segregating recessive deleterious alleles, this is supported by the absence of inbreeding depression and evidence for outbreeding depression (lower hatching success when individuals from isolated populations were mated) in one of the social species *S. dumicola* (Berger-Tal et al. 2014).

**Figure 1.**
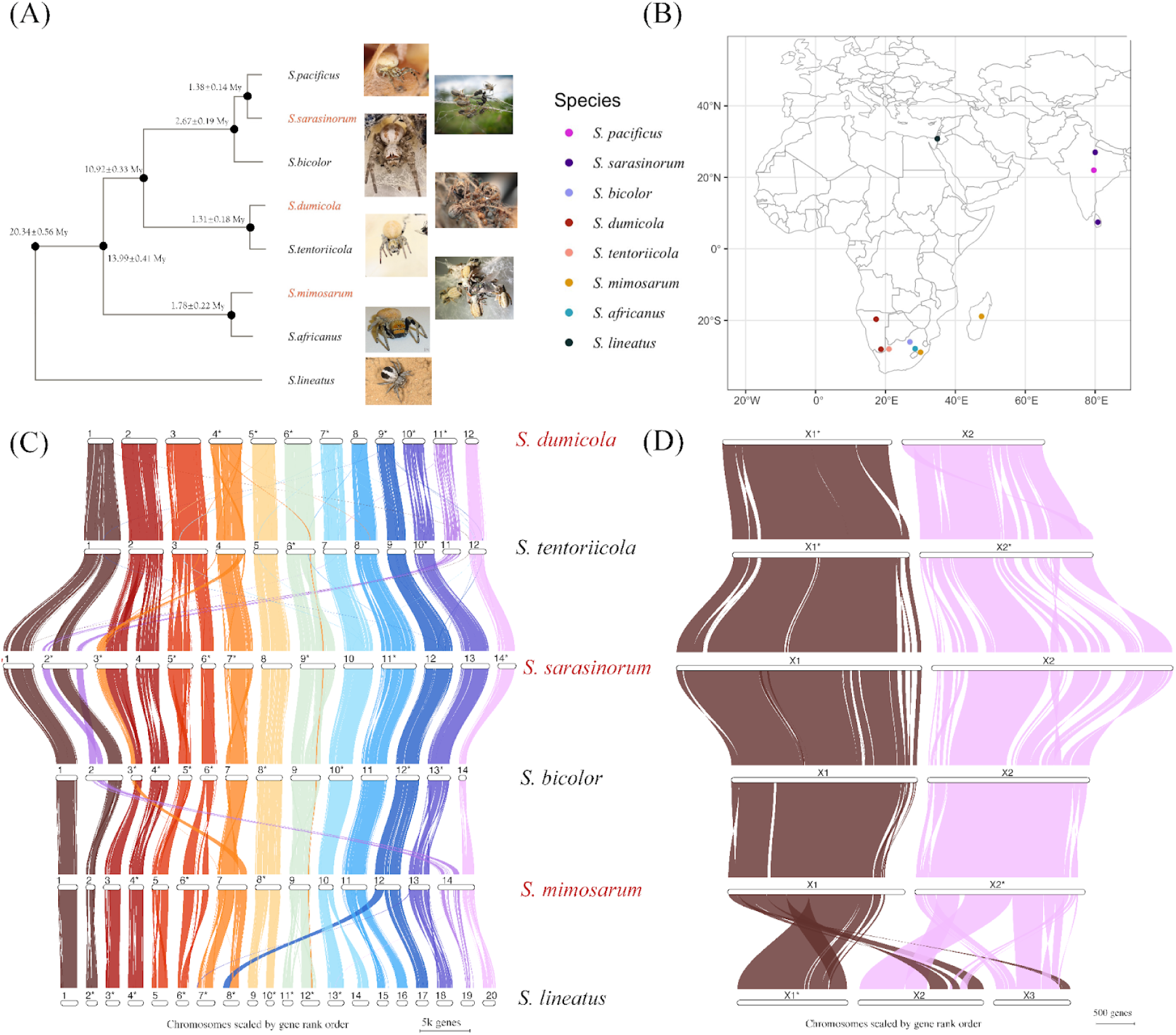
(A) Phylogeny of the spider genus *Stegodpyhus* with social species in red. The divergence time is estimated from *d*_S_ of autosomal single-copy orthologous genes and a yearly mutation rate of 5e-09 per site per year. (B) Sampling locations of the species included in our study. The gene synteny plots among the six *de novo* chromosome-level assemblies are shown using GENESPACE, separately for autosomes (C) and X chromosomes (D). The length of chromosomes in the gene synteny plot represents the number of genes. The id of chromosomes in (C) and (D) are labelled as the number we assigned in the assembled genome of each species. Stars(*) mark chromosomes that are complementary-reversed in the synteny analysis.

Here we performed whole genome assembly and comprehensive comparative analyses of the three social *Stegodyphus* species and their subsocial sister-species to assess the timing of evolutionary transitions and genomic consequences of social evolution. Based on analyses of Ne trajectories and purifying selection, we determined the divergence time of lineages and the age of individual transitions to sociality, to assess whether speciation coincides with the evolution of sociality. This information further serves to investigate the progression of genomic consequences caused by the transition to sociality via comparisons of social spider lineages of different ages. These data sets enable us to assess the hypothesis that transitions to sociality are evolutionary dead-ends in spiders.

## Results

### *De novo* assembly and genome annotation

Six species of *Stegodyphus* (3 social, 3 subsocial) were sequenced to ∼30X coverage using Pacbio HiFi long reads. Combining this with Hi-C contact maps based on 30X Illumina sequencing, we generated chromosome-level, *de novo* assemblies. (Table 1, Supplementary Figure S1). The sex chromosomes of each species were identified by the read coverage of male individual alignments. We find two sex chromosomes in all species except for the outgroup species *S. lineatus* which has three X chromosomes (Figure S8). For each species, the inferred number of linkage groups matched the number of chromosomes found by cytological visualisation (Avilés et al. 1999; Král et al. 2011). We evaluated possible assembly errors by comparing the independent assemblies by dot plots using dotPlotly (Supplementary Figure S4-S6). For the most closely related species pairs (1-3% divergent at the genome level), assemblies were essentially syntenic, suggesting that the rate of assembly error is very low and also that the synteny breaks for more divergent species pairs indicate real chromosomal rearrangements and/or inversions. Annotation using BRAKER2 (Brůna et al. 2021) independently for each assembly revealed similar numbers of protein-coding genes ranging from 32,093 to 35,724, except for *S. sarasinorum* where 47,066 protein-coding genes were annotated. This recent gene number expansion in *S. sarisinorum* is not an artefact of annotation and is the focus of a separate study. BUSCO (Simão et al. 2015) analyses show high completeness of the genomes ranging from 91.1% to 93.3%. The GC content ranges from 33.26% to 34.18%, and the proportion of the genomes constituting repeats ranges from 57.17% to 65.20%. We found 10,065 single-copy orthologous genes present in all six species using our *de novo* assemblies, synteny information and annotations (Table 1, Table S2). Mercury (Rhie et al. 2020) estimation on our genomes reveal k-mer completeness ranging from 97.65% to 99.41% and a consensus quality QV ranging from 62.54 to 73.86, which is higher than the Vertebrate Genomes Project (VGP) benchmarking recommendations of QV > 40 and k-mer completeness > 95% (Rhie et al. 2021). When incorporating the *de novo* assembled transcriptome from *S. africanus* and short reads DNA sequences from *S. pacificus*, our final dataset ends up with 2,303 and 348 reliable single copy orthologs for autosomes and X chromosomes, respectively, across all 8 species.

**Table 1.** Assembly and annotation statistics. BUSCO completeness is only reported for the single complete fraction checked with protein sequences from annotated genes, complete BUSCO results reported in Table S2.

| Species | Chrom_Num | Sex_chrom | Scaffolded Length | GC level | RepeatMasked | Gene_Num | BUSCO (S.C) |
| --- | --- | --- | --- | --- | --- | --- | --- |
| *S.lineatus* | 23 | X1X2X3O | 2658234335 | 33.26% | 58.72% | 32093 | 94.10% |
| *S.mimosarum* | 16 | X1X2O | 2910110412 | 33.92% | 57.17% | 35480 | 92.40% |
| *S.tentoriicola* | 14 | X1X2O | 2984342755 | 33.42% | 64.59% | 35725 | 91.30% |
| *S.dumicola* | 14 | X1X2O | 3146466829 | 34.18% | 64.53% | 34096 | 91.00% |
| *S.bicolor* | 16 | X1X2O | 2984548140 | 33.78% | 61.98% | 35096 | 92.30% |
| *S.sarasinorum* | 16 | X1X2O | 3321123892 | 34.06% | 65.20% | 47066 | 93.10% |

### High gene synteny and few chromosomal rearrangements in the *Stegodyphus* genus

To investigate genome rearrangements across *Stegodyphus* from the accurate *de novo* assemblies, we used GENESPACE (Lovell et al. 2022) to retrieve the pair-wise synteny blocks based on the gene order of orthologs found in each *de novo* annotation (Figure 1 outlines the results). In general, synteny blocks suggest few chromosomal translocations and further support that the assemblies are accurate.

We identify a number of large-scale structural rearrangements including splits and joins of chromosomes unique to certain lineages of the phylogeny. For example, the outgroup species *S. lineatus* has 23 chromosomes including three X chromosomes, while the other species contain 14 to 16 chromosomes and only two X chromosomes, suggesting that multiple chromosome splits or joins have happened in the *S. lineatus* lineage, or that chromosome joins have occurred after the split with *S. lineatus* in the common ancestor of the other species (Table 1). We find that the third X chromosome in *S. lineatus* appears to originate from a “cut and paste” non-homologous recombination process, which joins a part of the ancestral X chromosome X1 and a part of the ancestral X chromosome X2 (Fig 1. D). Thus, the three X chromosomes in the *S. lineatus* lineage have almost the same gene complement as the two X chromosomes in the other species. In addition, we find three chromosome join events assigned to the ancestral branch of *S. dumicola* and *S. tentoriicol* (chromosome 1, 2, and 3 are homologous to 6 separate chromosomes in other species), and one chromosome join event (chromosome 2 are homologous to 2 separate chromosomes in other species) and a non-homologous recombination event (chromosome 3 contain extra components that are homologous to another chromosome in other species) assigned to the ancestral branch of *S. bicolor* and *S. sarasinorum*.

### Estimating the beginning and end of the transitions to sociality

We first estimated divergence times at each node of the *Stegodyphus* phylogeny using the resampled branch-wise *d*_S_ from autosomal branches and assuming a mutation rate of 5e-09 per site per generation (Wang and Obbard 2023). Among the three pairs of social and subsocial sister species, the estimated divergence times are 1.78±0.22 million years between *S. mimosarum* and *S. africanus*, 1.31±0.18 million years between *S. dumicola* and *S. tentoriicola*, and 1.38±0.14 million years between *S. sarasinorum* and *S. pacificus*.

Transitions to sociality in spiders are associated with a shift to inbreeding and reduced effective population size (Ne) causing natural selection to be less efficient. We assume that a substantial less efficient selection and Ne reduction in the social spider species are more likely caused by evolution of sociality than other factors. We then used our genome data to estimate the time when these transitions to sociality occurred from the expected signals in purifying selection and effective population size (Ne). We did this in two separate ways, first from the estimated changes in purifying selection estimated from *d*_N_/*d*_S_ ratios, and second, from the inferred changes in effective population size through time estimated from coalescent analyses.

#### Intensity of purifying selection

We estimated the transition time to social behaviour from *d*_N_/*d*_S_ ratios on the terminal branches of each social/subsocial species pair based on the following assumptions: a) *d*_N_/*d*_S_ ratios of the social species was the same as for the subsocial species until social transitions on the terminal branches leading to the social species, b) after transition to sociality, the *d*_N_/*d*_S_ ratio increased due to less efficient purifying selection, c) this ‘social *d*_N_/*d*_S_’, can be approximated by the *pi*_N_/*pi*_S_ ratio estimated between divergent and isolated extant populations of each social species (see Methods). This *d*_N_/*d*_S_ analysis suggests that the effects of a transition to sociality on purifying selection are very recent (Figure 2 A-D). For example, the *d*_N_/*d*_S_ over the entire *S. mimosarum* branch is 0.1259±0.0044, and for its subsocial sister-species *S. africanus d*_N_/*d*_S_ is 0.0927±0.0007. The *d*_N_/*d*_S_ ratio since sociality evolved is estimated at 0.3043±0.0018 comparing two *S. mimosarum* individuals from Madagascar and mainland Africa. From the divergence time estimates for this species pair, this generates an estimated time since transition to sociality of *S. mimosarum* of 279 kya (95% CI: 230 kya - 332 kya). Using the same calculation, the social *d*_N_/*d*_S_ ratio of *S.dumicola* is estimated to be 0.3366±0.0037 by comparing individuals from Otavi North Namibia and Karasburg South Namibia. Combining the *d*_N_/*d*_S_ ratio being 0.1776±0.0039 and 0.1615±0.0054 for the *S. dumicola* and *S. tentoriicola* branches respectively, the estimation of the transition to socialility is 119 kya (95% CI: 72 kya - 170 kya) in *S. dumicola*.

**Figure 2.**
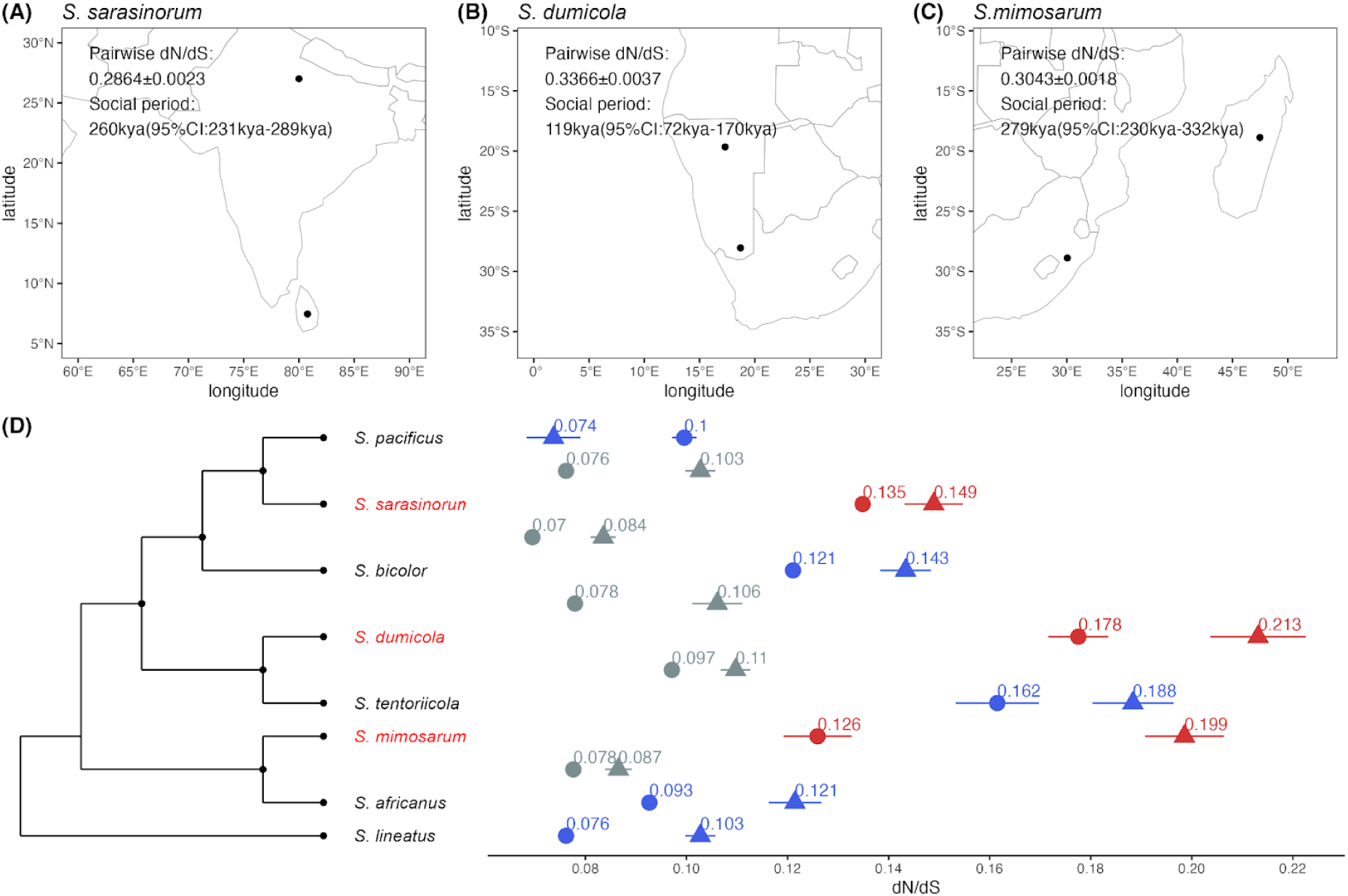
The intensity of purifying selection was assessed using *d*_N_/*d*_S_ from the coding sequences of 2302 single-copy orthologous genes. (A) The locations of the Sri Lanka *S.sarasinorum* and Himalayas *S.sarasinorum* samples, along with their pairwise *d*_N_/*d*_S_ estimations for the social period. (B) The locations of the Namibia South *S.dumicola* and Namibia North *S.dumicola* samples, with corresponding pairwise *d*_N_/*d*_S_ estimations for the social period. (C) The locations of the South Africa *S.mimosarum* and Madagascar *S.mimosarum* samples, and their pairwise *d*_N_/*d*_S_ estimations for the social period. (D) Branch-wise *d*_N_/*d*_S_ estimations using PAML, with the 95% confidence interval from resampling. Red, blue, and grey denote social species terminal branches, subsocial species terminal branches, and internal branches, respectively. The *d*_N_/*d*_S_ estimation, indicated by points with round and triangle denote autosomes and X chromosomes respectively, is aligned to the phylogeny that corresponds to the respective branches. The phylogeny in (D) provides an indication of the topology without branch length information.

For the social *S. sarasinorum,* we performed two separate estimations by comparing *S. sarasinorum* to the subsocial species *S. pacificus,* which diverged 1.38±0.14 million years ago, and the subsocial species *S. bicolor*, which diverged 2.67±0.19 million years ago. In *S. sarasinorum*, we estimated the social *d*_N_/*d*_S_ ratio to be 0.2864±0.0023 based on two individuals from Himalaya and Sri Lanka. The *d*_N_/*d*_S_ ratio on the terminal branch of *S. sarasinorm* is 0.1349±0.0009. Compared to the 0.0996±0.0016 of the *S. pacificus* branch and the 0.1211±0.0007 of the *S. bicolor* branch, the final estimation for the transition to sociality of *S. sarasinorum* is 260 kya (95% CI: 231 kya - 289 kya) and 223 kya (95% CI: 199 kya - 247 kya), respectively.

We have attributed the observed higher *d*_N_/*d*_S_ ratio in the terminal branches of social species to a decrease in purifying selection. However, increased adaptive selection acting on social species could have the same effect. RELAX analyses (Wertheim et al. 2015) suggest that the elevated *d*_N_/*d*_S_ ratios in social species is almost entirely due to relaxed selection in autosomes (Figure S3), as has previously been suggested (Tong et al. 2022). We also show elevated *d*_N_/*d*_S_ ratios on X chromosomes compared to autosomes across all branches in the phylogeny, except for the terminal branches of *S. pacificus*.

Such elevated *d*_N_/*d*_S_ ratios can be attributed to a decrease in purifying selection acting on X chromosomes, which is consistent with the expected outcome of reduced Ne in X chromosomes compared to autosomes (Ne_X/Ne_Auto is expected to be 0.75 in diploid species with equal sex ratio). Alternatively, but not mutually exclusive, an adaptive explanation for elevated *d*_N_/*d*_S_ ratios on the X chromosomes is the so-called faster-X effect (Charlesworth et al. 1987). Recessive variants on the X chromosomes are directly exposed to selection in males that are haploid for X chromosomes, and under certain conditions this will lead to more adaptive substitutions on X chromosomes compared to autosomes.

Additionally, we note that our implication of using *pi*_N_/*pi*_S_ between divergent populations of the same species could potentially lead to a systematically higher “social *d*_N_/*d*_S_” due to the unfixed polymorphisms rather than reflect the actual selection strength when the selection strength acts differently on synonymous polymorphism sites and non-synonymous polymorphism sites within species. We benchmarked our strategy by comparing the site frequency spectrum of missense variants (N-sites) and synonymous variants (S-sites) in *S. dumicola* built from 9 individuals. Site frequency spectrums of the N-sites and S-sites reveal no visually significant difference in shape, indicating genetic drift being the major force acting on both N-sites and S-sites (Figure S7). Mattila et al 2012 also quantified the *d*_N_/*d*_S_ and *pi*_N_/*pi*_S_ in three *Stegodyphus* species, including one social species (*S. mimosarum)* and two subsocial species (*S. tentoriicola* and *S. lineatus*) (Mattila et al. 2012). Mattila et al. reveal that the increase of *pi*_N_/*pi*_S_compared to *d*_N_/*d*_S_ is much higher in the social spider species (3-fold) than subsocial species (1-1.5 fold)(Mattila et al. 2012). It is also worth noting that the *pi*_N_/*pi*_S_ in Mattila et al 2012 is based on transcriptome data from a single individual, where the unfixed polymorphisms will increase the tendency of inherently higher *pi*_N_/*pi*_S_ than *d*_N_/*d*_S_ at a larger effect than using isolated populations. Thus, we conclude that *pi*_N_/*pi*_S_ to a good approximation can be used as a proxy for the *d*_N_/*d*_S_ ratio after sociality (“social *d*_N_/*d*_S_”).

#### Coalescent analysis

Next, we used PSMC (Li and Durbin 2011) analysis to reconstruct the effective population sizes and the sex ratio in the populations over time by analysing autosomes and X chromosomes separately in each species. For the social species, which are all highly inbred, a diploid genome was created by merging two haploid genomes sampled from different divergent and isolated populations, i.e. an artificial diploid. This allows coalescent events further back in time to be inferred. The transition time to sociality is then estimated by graphically determining the time period when the effective population size of the social species became severely reduced (Figure 3). Compared to subsocial species, the population trajectories of all the social species share a gradual reduction of Ne from more than 300,000 to less than 50,000, but starting and ending at different time points within the past 900 ky. In the social species *S. dumicola*, the Ne is reduced from 300,000 around 420 kya to around 50,000 at 300 kya, and even lower today. In the social species *S. mimosarum*, the Ne is reduced from 1.4 million to 57,500 in the time interval between 900kya and 350kya. The reduction of Ne in the social species *S. sarasinorum* starts 600 kya at 800,000 to less than 30,000 around 300 kya.

**Figure 3.**
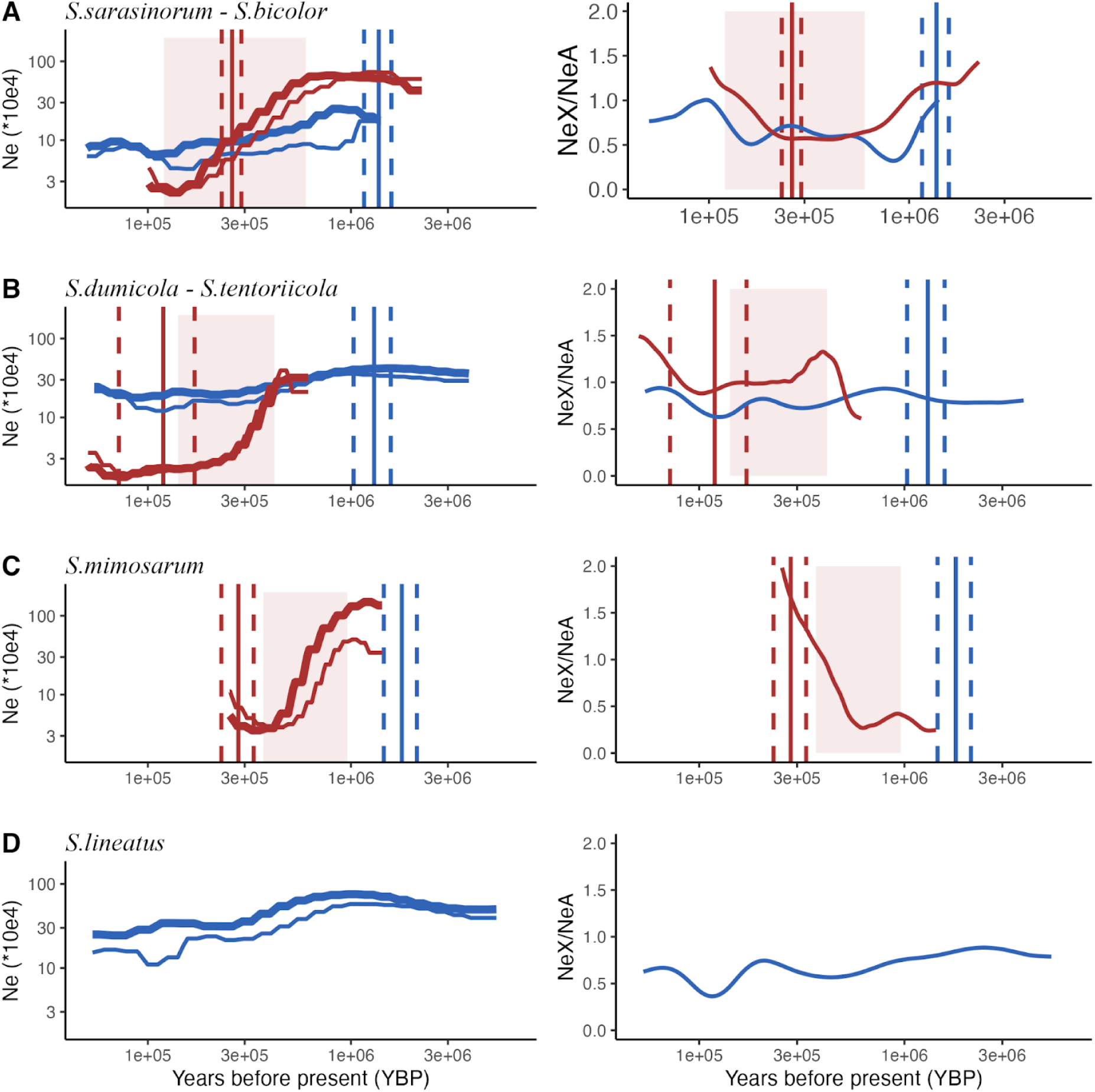
The historical effective population size inferred from the Pairwise Sequentially Markovian Coalescent (PSMC) model, for different *Stegodyphus* species with chromosome-level assembly. In each panel, the left figure depicts PSMC results, while the right figure displays the effective population size ratio between X chromosomes and autosomes. Each species’ trajectory is shown with thicker lines for autosomes and thinner lines for X chromosomes. The type of species are distinguished by colour: red for social species and blue for subsocial species. Red shadows mark the estimated social transition period based on the decreasing effective population size in social species. Vertical lines indicate estimated time points by comparing the social species with its closest sister subsocial counterpart. Red lines correspond to the estimated social transition time, derived from relaxed selection in social species, while blue lines represent the estimated divergence time between a social species and its closest subsocial sister species. Solid lines provide point estimates, and dashed lines delineate the 95% confidence intervals. The PSMC results are truncated in the near recent past for better comparison across species. Bootstrapped PSMC results without time truncation are provided in Figure S9. (A) Displays data for the social species *S.sarasinorum* and the subsocial species *S.bicolor*. Estimates for social transition and divergence time are based on a comparison between *S.sarasinorum* and its subsocial sister species, *S.pacificus*. (B) Presents findings from a comparison between the social species *S.dumicola* and its subsocial sister species *S.tentoriicola*. (C) Shows data for the social species *S.mimosarum*, with social transition and divergence time estimates based on a comparison with its subsocial sister species, *S. africanus*. Note that we do not have an effective population size estimation for *S. africanus* due to missing whole genome data. (D) Presents historical effective population size for the outgroup subsocial species *S.lineatus*.

Since the PSMC analyses of the social species are made from pseudodiploid genomes combined from separate genetically isolated populations, they can additionally provide an estimate of the time of separation of these populations, since coalescence can not occur when they are separate and the Ne estimate will thus be very large (Figure S9). We observe such an uptick in Ne in recent times in the three social species, and estimate that the two *S. mimosarum* populations from Madagascar and South Africa diverged around 200 kya, the two *S. dumicola* populations in North and South Namibia diverged 20 kya, and the two *S. sarasinorum* populations from Himalaya and Sri Lanka diverged approximately 110 ky ago.

The pseudo-diploid of social species collapsed two heavily inbred diploid individuals. We estimate the run out of heterozygosity (ROH) regions takes 59.83%-86.27% of their genomes (Table S4), but the individuals are still not 100% homozygous. Hence, they are not identical as a haploid in practice. Thus, we can not rule out the fact that pseudo-diploid actually capture more than two haploids for part of the genomes outside the ROH region. However, we expect this will only increase a local coalescence rate that is randomly distributed in the genome, resulting in a lower estimation of Ne overall, without biassing the Ne reduction information retrieved from the Ne trajectories.

### Evolution of sex ratio bias

The sex determination system of spiders is X0, and as there are four autosomes for each three X chromosomes in the population with 1-to-1 sex ratio, the effective population size of X chromosomes is expected to be 75% of the autosomes. In subsocial species, PSMC analyses of X chromosomes and autosomes showed that although the estimated Ne for X chromosomes is indeed lower, the ratio of X chromosomes to autosome effective population size is around 50-60% instead of the expected 75%. This can be due to a lower mutation rate and/or stronger selection on the X chromosomes, as was previously suggested based on diversity estimates on X chromosomes and autosomes in *S. africanus (Bechsgaard et al. 2019)*.

In contrast to the subsocial species, the social species show strongly female biassed primary sex ratios (Lubin and Bilde 2007) and this has been shown to be genetically determined (Vanthournout et al. 2018). When a species evolves from an unbiased to a female biassed sex ratio, the effective population size of X chromosomes should approach that of autosomes. In concordance with this, the PSMC analyses reveal that X chromosome and autosome Ne are almost identical in the most recent time intervals of the PSMC, as expected when sociality has fully evolved. The evolution of a female biassed sex ratio can therefore also be timed using PSMC analyses under the assumption that it occurred after the transition to sociality, by comparing the Ne of X chromosomes and autosomes back in time. We found elevated Ne(X)/Ne(autosomes) ratios in all social species associated with reductions in Ne, which links the evolution of sex-ratio bias with social transitions. This suggests that the evolution of a female biassed sex ratio in the three social species occurred early in the evolutionary transition to sociality.

The evolution of female bias is predicted under strong continuous inbreeding and lack of pre-mating dispersal, characteristics of social spider species and other social arthropods (Johnson et al. 2000; Chapman et al. 2000; Keller et al. 2011), to reduce competition among brothers for fertilisation of their sisters (Bulmer and Taylor 1980; West 2009). Furthermore, allocation to daughters as the helping sex in social species is expected by the local resource enhancement model (Trivers and Willard 1973). Biassed sex ratios are often caused by sex chromosome meiotic drive (Lindholm et al. 2016), and the driving loci are selfish genetic elements that would usually trigger adaptations to restore even sex ratio (Lindholm et al. 2016). However, selection against female bias is not expected here, because of the adaptive benefits of female biassed sex allocation (Vanthournout et al. 2018) generating predictions of rapid fixation of the sex-linked meiotic driver. If stronger female bias keeps being adaptive, selection will lead to an increasingly female biassed sex ratio following transition to sociality. The results from the PSMC analyses showing that NeX/NeA continues to increase after transition to sociality suggest that stronger female bias continues to be adaptive. Furthermore, our estimation for the transition to sociality matches with observations of increasing female bias with age of sociality in the three species. The oldest social species *S. mimosarum* has the strongest sex ratio bias with 9% males, the younger social species *S. dumicola* and *S. sarasinorum* have a male percentage of 12% and 8.5-33%, respectively (Lubin and Bilde 2007). The correlation between sex ratio bias and age of the social period in the three transitions to sociality therefore indicate that the continued loss of males is genetically determined.

## Discussion

### Divergence times and transition to sociality

To elucidate genomic consequences of repeated, independent transitions to sociality, we estimated species divergence times and the time of speciation using different approaches (Figure 4). An obvious question is whether the speciation itself was caused by one of two diverging populations evolving sociality. The large time interval between estimates of species divergence (Td = 1.3-1.8 my, Figure 4) and the beginning of social transitions (400-900 ky) as derived from strong reductions in Ne at first suggests that species separation took place first with subsequent evolution of sociality and cooperative breeding. However, coalescent theory predicts that it takes 2Ne generations on average for ancestral polymorphisms to find most common ancestry. Since the effective population sizes of subsocial species are very large (300,000-600,000), divergence times (Td in Figure 4) are expected to be much more ancient than the species separation times (Ts in Figure 4) due to long coalescence times in the ancestral species. With a generation time of one year, average coalescence time in the ancestral species are between 600,000 and 1.2 million years, and our best estimate of the speciation times (Ts in Figure 4) are therefore almost 1 million years closer to the present than the divergence times. This corresponds to the time when we estimate the beginning of the social transitions to (from PSMC results), thus suggesting that social behaviour emerging in a population of a species may facilitate separation of this population into a new species.

**Figure 4.**
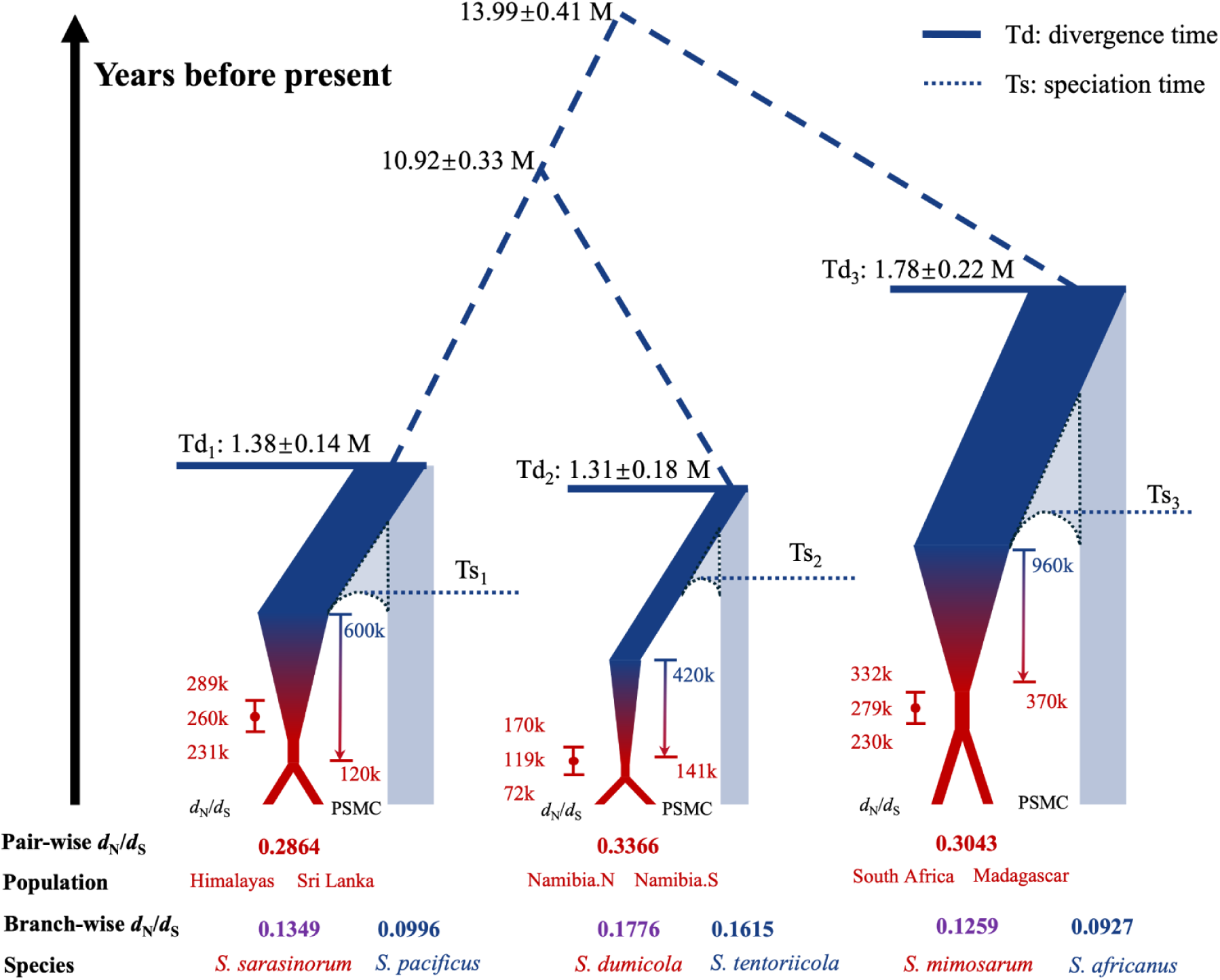
Estimated species divergence (Td), and the time of transition to sociality/cooperative breeding (indicated in red) in three independent social transitions within the *Stegodyphus* spider genus. Hypothetical speciation time (Ts) between each species pair is marked as the dashed lines. The scale of the plots is symbolic and not proportional. The widths of the terminal branches represent variation in effective population size, with red symbolising social periods and blue indicating subsocial periods. Light blue branches represent species without chromosome-level assemblies for PSMC. For each lineage where a social transition occurs, point estimates and 95% confidence intervals for social transition times are presented on the left, while social transition periods derived from population size history are displayed on the right. In PSMC estimations, the start of social transition periods corresponds to the initial decrease in effective population size, while the end coincides with Ne dropping below 50,000.

There is, however, also some evidence from the social *S. dumicola* that sociality can arise independently after the speciation process. *Stegodyphus dumicola* and *S. sarasinorum* diverged from their closest subsocial sister species at a similar time scale of 1.30 Mya and 1.37 Mya respectively. The ancestral Ne for *S. dumicola* (300,000) is smaller than the *S.sarasinorum* (600,000). Thus, the speciation time of *S. dumicola* (Ts2 in Figure 4) is expected to be closer to the species divergence time than the speciation time of *S. sarsinroum* (Ts1 in Figure 4) based on coalescence theory (Figure S11). Meanwhile, both analysis from PSMC and *d*_N_/*d*_S_ show that the age of sociality is younger in *S. dumicola* than *S. sarasinorum*. The combination of the earlier speciation time and the younger age of sociality in *S. dumicola* compared to *S. sarasinorum* suggests that sociality can arise as a separate event from the initiation of speciation. Nevertheless, evidence for separation of the two events in *S. dumicola* could be confounded by other factors, including (a) a missing subsocial sister species (unidentified or extinct) to *S. dumicola* that is not included in the analysis, which can bias the analyses to overestimate the divergence time for *S. dumicola* lineage, (b) observed reduction of Ne in recent past of *S. tentoriicola* from PSMC (Figure S9) and elevated *d*_N_/*d*_S_ in *S. tentoriicola* compared to other subsocial species (Figure 2), which can bias the analyses to underestimate the age of sociality in *S. dumicola*. Nevertheless, the combination of data presented above substantiates that sociality and characteristics of the ‘social syndrome’ evolved much more recently than the divergence estimate.

### High extinction risk of social spiders

The genomic consequences of transitions to sociality in spiders are consistent with expectations of strong reductions in Ne when the evolution of sociality is accompanied by a transition from outcrossing to inbreeding and the development of a female biased sex ratio (Vanthournout et al. 2018). The relatively young age of social lineages indicate that sociality evolves rapidly and frequently in the spider phylogeny and comes with a subsequent strong signal of decreased purifying selection caused by strict inbreeding and much reduced effective population size. Overall, there are at least 23 independent convergent origins of sociality (Lubin and Bilde 2007; Agnarsson et al. 2006). The recurring and rapid evolution of cooperative breeding, coupled with the absence of lineages with a long history of sociality, is strong indirect evidence that sociality in spiders represents an evolutionary impasse, unlikely to persist beyond a timespan of 1 millions years once established. Thus, it is plausible that numerous social spider species have emerged and gone extinct throughout evolutionary history, but we can only observe the presence of extant and recently developed social spider species.

The risk of extinction in social spiders is hypothesised to be much higher than in eusocial insects owing to the ‘social syndrome’ which causes reductions in Ne. Phylogenetic studies show that diversification of species is only associated with a few old transitions to eusociality in the order *Hymenoptera*, i.e. ants, bees and wasps(Wcislo and Fewell 2017; Hunt and Toth 2017; Heinze et al. 2017), which is different from the repeated and recent convergent evolution of sociality in the spider phylogeny (Agnarsson et al. 2006; Settepani et al. 2016). We propose that a key factor contributing to the difference in longevity of social lineages among major taxa is the elimination of pre-mating dispersal which results in an obligatory inbreeding mating system. In eusocial insects, new colonies are established by mated queens, and outcrossing is maintained by pre-mating dispersal of reproductive individuals (Wcislo and Fewell 2017; Heinze et al. 2017). Thus, eusocial insects may show reduced Ne and relaxed selection compared to their nonsocial counterparts (Kapheim et al. 2015), but the reduction in Ne reduction is mitigated by the maintenance of outcrossing. In contrast, we here show that transitions to sociality in independently evolved spider lineages are associated with dramatic declines in Ne.

### Chromosomal systems, mating systems and the evolutionary longevity of social lineages

Some of the most successful social lineages have evolved in the *Hymenoptera*, where the entire clade is haplodiploid with haploid males and diploid females (Grimaldi and Engel 2005). These lineages are evolutionarily old, for example in ants where eusociality is dated at least to the Cretaceous (Table 2) and they have since diversified dramatically both in species and abundance to become one of the most ecologically dominating taxonomic groups in the world (Wilson and Hölldobler 2005). The evolutionary success of haplodiploid social lineages, which is also found in *Thysanoptera* (thrips) and sporadically in some spider mites, *Hemiptera*, and *Coleoptera*, has long pointed to an importance of haplodiploidy in the evolution of sociality (Hamilton 1964; Trivers and Hare 1976; Joshi and Wiens 2023). However, phylogenetic patterns are debated and perhaps inconclusive, as eusociality has evolved repeatedly within the haplodiploid *Hymenoptera* and *Thysanoptera*, whereas other successful independently evolved social lineages are diploid, such as in termites and aphids.

**Table 2.** Summary of the characteristics of sociality in diploid and haplodiploid organisms with different mating systems. *The social complexity is a broad qualitative summary with the terminology used for describing social diversity in insects in Fewell and Abbot 2018.

| Ploidy | Mating system | Age of social lineages | Social Complexity*(F ewell and Abbot 2018) | Taxonomy range | Taxa examples |
| --- | --- | --- | --- | --- | --- |
| Diploid | Strictly inbreeding (sib-mating) | Young (Sociality only has full impact on selection within the past 300ky from this study) | Cooperative communal groups | Broad, often multiphyletic for the young social species lineages | Spiders(Agnars son et al. 2006; Settepani et al. 2016) |
|  | Outcrossing | Various? (oldest can be 130 million year in termites(Krishna et al. 2013)) | Advanced eusocial, Primitive eusocial, Cooperative communal groups |  | Termites(Legendre et al. 2008; Korb and Thorne 2017), Snapping shrimps(Chak and Rubenstein 2019), Mole rats(Kverková et al. 2018; Faulkes and Bennett 2021), Aphids(Stern 1994) |
| Haplodiploid | Inbreeding (sib-mating), occasionally outcrossing | Less Old? (10 million years in thrips, in term of species lineage(Abbot and Chapman 2017)) | Primitive eusocial, Cooperative communal groups | Limited (order *Hymenoptera* and *Thysanoptera*, tribe *Xyleborini*), often monophyletic for the old social species lineage. | Thrips(Chapman et al. 2000), Beetles(Johnson et al. 2018) |
|  | Outcrossing | Old (The phylogeny of ants dates back to 140 - 168 million years ago(Moreau et al. 2006)) | Advanced eusocial, Primitive eusocial |  | Ants, bees, wasps |

A key question that arises from our study is whether haplodiploidy represents a more robust system for governing the evolutionary longevity of social lineages. Termites and *Hymenoptera* represent the oldest social lineages and represent both chromosomal systems (Table 2). These taxa also maintain predominantly outcrossing mating systems, which acts to maintain higher effective population sizes that are less exposed to genetic drift. In taxa where social evolution is accompanied by a transition to inbreeding, the diploid social spiders show the most short-lived and recent transitions (Settepani et al. 2017), whereas thrips and beetles with a haplodiploid system may have relatively older social lineages (Table 2). In combination, these patterns suggest that whether or not sociality is associated with a transition from outcrossing to inbreeding is perhaps the most important factor for determining their evolutionary longevity. The taxa that evolve inbreeding mating systems are simultaneously characterised by female biased sex ratio which further acts to reduce effective population size (Bechsgaard et al. 2019). Our study provides genomic support for the hypothesis that the independent evolution of sociality accompanied by an inbreeding mating system and female-biased sex ratios results in drastic reductions in Ne and relaxed efficacy of selection, which likely contribute to drive lineage extinctions.

### Genomics determinants of evolutionary dead ends in social spiders

We note that even though the three social transitions we report were initiated at different times, they have followed remarkably similar trajectories in population size reduction, increased *d*_N_/*d*_S_, and biased sex ratios that complements the strong similarities in behaviour and life history associated with transitions to sociality. This almost deterministic genomic response to sociality may also hold the key to understanding why social spider species do not seem to persist for very long. Several possibilities may explain the demise. These include a) loss of adaptive potential due to loss of genetic diversity, b) fixation of deleterious variation, c) loss of males, and d) loss of genome integrity. While a) and b) may definitely make sociality a doomed evolutionary strategy, it is questinable whether these processes can happen within a few hundred thousand years since the fixation of slightly deleterious variants occurs over a very large number of generations (Lynch et al. 1995). Loss of males could carry on until population collapse, and it is intriguing that we see fewer males in the social transition estimated to be the oldest (Figure 4). Finally, loss of genome integrity by, for example transposable elements (TEs), increased mutation rate, poorer genomic repair of damage, and double strand breaks, are all global mechanisms that could impact the longevity of the social species. TEs are expected to accumulate in species with reduced Ne. Although the question of whether the TE fraction increases with reduced Ne remains inconsistent in several tested organisms (Glémin et al. 2019), evidence for increased amounts of TEs has been seen in eusocial snapping shrimps with reduced Ne (Chak et al. 2021). This idea warrants future research in social spiders.

## Methods

### Sample collection across the *Stegodyphu*s genus

The spider genus *Stegodyphus* is distributed widely across Eurasia and Africa and we collected seven species including three social and four subsocial species in this study (Supplementary Table 1). Samples were collected from multiple regions for social species (Fig. 1 A). We collected *S. dumicola* samples from Otavi and Betta in Namibia (2017), *S. mimosarum* samples from Antanario in Madagascar and Weenen in South Africa (2011), and *S. sarasinorum* samples from the Himalaya region and Sri Lanka (2014). For subsocial species, we collected *S.lineatus* from the Negev Desert in Israel (2020), *S. bicolor* from Betta in Namibia (2021), *S. tentoriicola* from Tierberg in South Africa (2015). *Stegodyphus africanus* RNA data was previously published in Bechsgaard et al (2019).

### Reference genome: DNA extraction, sequencing and assembly

DNA was extracted from single individuals using the MagAttract HWM DNA kit from Qiagen (Hilden, Germany). Sequencing was done by PacBio HiFi technology to obtain from 69 to 99 GB data.We assembled haplotype resolved contigs using Hifiasm(Cheng et al. 2021) with Pacbio HiFi reads and Hi-C reads. Hi-C reads were aligned to the contigs from the longer haplotype of each species using Juicer pipeline for scaffolding chromosome-level assemblies using 3D-DNA pipeline (Dudchenko et al. 2017). The chromosome-level assemblies were further manually checked and curated using Juicebox Assembly Tools(Durand et al. 2016) based on consistency of Hi-C contact pattern.

### Resequencing: DNA extraction and sequencing

DNA was extracted from single individuals using the Qiagen Blood and Tissue kit, Qiagen (Hilden, Germany), and the DNA was sequenced using DNBSEQ-G400.

### Genome Annotation

We annotated six species with chromosome-level assemblies using BRAKER2 with RNA-seq and protein homology evidence combined. For each species, we used RepeatModeler2(Flynn et al. 2020) to generate a repeat library and combine it with the repeat library for Arthopoda from Repbase(Bao et al. 2015). The chromosome-level assemblies are masked using Repeatmasker(Tarailo-Graovac and Chen 2009) and their corresponding repeat libraries. We used STAR (Dobin et al. 2013) to align RNA-seq reads to the repeat-masked genome as transcriptome evidence for gene prediction in BRAKER2. For protein homology evidence, we used protein sequence from a previous *S.dumicola* annotation (Liu et al. 2019) and arthropoda protein from OrthoDB v10 (Kriventseva et al. 2019). In the end, the completeness of the genome annotations is evaluated with BUSCO together with a set of 1013 *Arthropoda* genes from OrthoDB v10.

### Ortholog groups finding and multiple genome alignments

We performed GENESPACE to check the gene synteny among the six hifi chromosome-level assemblies. During the GENESPACE analysis, OrthoFinder (Emms and Kelly 2019) was applied to identify orthologous groups across the six species. The 10065 single-copy orthologous groups out of 48143 orthologous groups identified from the six species were further used in evolutionary analysis.

We do not have the genome assembled from *S. africanus*. To include *S. africanus* in the analyses, we downloaded the transcriptome data of *S. africanus* from Bechsgaard et.al 2019 (Bechsgaard et al. 2019) and used Trinity (Haas et al. 2013) to assemble transcripts. We used DIAMOND(Buchfink et al. 2021) blastx mode to align the transcripts to the database of all the translated amino acid sequences from the 10065 single-copy orthologous groups of all six species. 5590 out of 10065 single-copy orthologous groups find one or more hits from the trinity transcriptome assemblies. We further check the aligned percentage of the hitted orthologous genes and filtered for a percentage of 75% to consider a transcript from *S. africanus* being a valid match to a certain single-copy ortholog groups identified, which ended up with 2649 single-copy orthologous groups.

We also do not have a genome assembled from *S. pacificus*. To include *S. pacificus* genes into the 2649 single-copy orthologous groups identified, we first used bwa-mem2 (Vasimuddin et al. 2019) to aligned short-read DNA sequence from a *S. pacificus* individual to the *S.sarasinorum* reference genome, which is the closest sister species of it. Then we used SAMtools (Danecek et al. 2021) consensus to call the consensus sequence as the genome of *S. pacificus* based on the bam file. The sequences of 2649 ortholog genes in *S. pacifcus* are then retrieved using the genome annotation file of *S. sarasinorum*. This process can be challenging when there are indels found in *S. pacificus* compared to *S. sarasinorum* reference when we construct the consensus sequence of *S. pacificus*. The sequence of *S. pacificus* and *S. sarasinorum* will not be in alignment as they have different total lengths with different genome coordinate systems. In practice, we fill gaps for deletions and remove insertions in *S. pacificus* based on the reference of *S. sarasinorum* when we construct the consensus sequences. This practice is achieved by setting parameters for SAMtools being “consensus --show-del yes --show-ins no”. By ignoring the indels in such a way, we maintain the same coordinate systems between the *S. pacificus* and *S. sarasinorum*. Orthologous sequences in S. pacificus can be then retrieved directly using the genome annotation file of *S. sarasinorum* to build multiple sequence alignments across all 8 species for *d*_N_/*d*_S_ estimations.

### Branch-wise *d*_N_/*d*_S_ estimation

We used CodeML in PAML(Yang 2007) to estimate *d*_N_ and *d*_S_ in branch-wise mode based on the 2649 single-copy orthologous groups. The 2649 orthologous groups were split up into an autosome set containing 2302 genes and a X chromosome set containing 347 genes. We randomly sample 500 genes from the 2302 autosomal without replacement in each resampling run, and we run the resampling independently 500 times to have an estimation of *d*_N_/*d*_S_ of each branch in the phylogeny with confidence intervals. For X chromosomes, we randomly sample 100 genes independently 100 times instead.

We do codon aware alignment in gene-wise for *d*_N_/*d*_S_ estimation. We used alignSequence from MACSE (Ranwez et al. 2018) to align coding sequences of each gene from all the species excluding *S.pacificus*, which was attached to the alignment according to the *S.sarasinorum* in the end. We also examined and filtered each gene alignment for local mis-alignment. The size of each alignment block without gap should have a length for at least 300 nucleotide base pairs as well as a fraction of polymorphism sites under 15% (Figure S2). The passing alignment blocks of the randomly chosen genes were concatenated using GOalign (Lemoine and Gascuel 2021) for *d*_N_/*d*_S_ estimation.

### *d*_N_/*d*_S_ (empirical *pi*_N_/*pi*_S_) estimation for social species

We used CodeML in PAML to estimate a pairwise *d*_N_/*d*_S_ for each social species with two separate populations. We aligned one individual from different populations to the species reference separately using BWA-MEM2. After calling the SNP using platypus (Rimmer et al. 2014), we retrieve the alternative nucleotide to substitute the species reference as the reference genome of each sub-population. Sequences from the 2649 single-copy orthologous genes were retrieved from the sub-population references for pairwise *d*_N_/*d*_S_ estimation. We used the same 500 resampled sets of genes as used in the branch-wise *d*_N_/*d*_S_ estimation before to estimate the *pi*_N_/*pi*_S_ with 95% confidence interval between separate social species populations for autosomes.

### *d*_N_/*d*_S_ and estimation of speciation divergence and social transition

For the divergence time between lineages, we used the dS branch length, a constant mutation rate of 5e-09 per site per generation, and a generation time of one year to scale the *d*_S_ length into years. We use the average *d*_S_ branch length between a split as an estimation of divergence time except for the split between *S.sarasinorum* and *S.pacificus*. The exception is due to the underestimation of *d*_S_ length of *S. pacificus* lineage from ignoring of indels, thus we use the *d*_S_ length of *S.sarasinorum* branch alone for divergence time estimation between *S.sarasinorum* and *S.pacificus*.

Assuming the transition to sociality is an instant process. The *d*_N_/*d*_S_ estimated for the social species terminal branch is an average of the subsocial period *d*_N_/*d*_S_ and social period *d*_N_/*d*_S_ weighted by the length of each period on that branch. The same idea has been applied to estimate the age of selfing in *Arabidopsis thaliana* compared to selfing incompatible *Arabidopsis lyrata (Bechsgaard et al. 2006)*. In our case, the subsocial period can be substituted approximately with the *d*_N_/*d*_S_ for the branch of the subsocial sister species. The social period *d*_N_/*d*_S_ can be estimated by pairwise comparing two individuals from two isolated and divergent populations of the social species.

For each resampling run of autosomal genes (500 genes out of 2303 genes), we obtained four estimations for each pair of social species and its subsocial counterpart using PAML, include

a. *d*_N_/*d*_S_ of the subsocial lineage
b. *d*_N_/*d*_S_ of the social lineage
c. *pi*_N_/*pi*_S_ between two isolated populations of social species
d. dS branch length between social and subsocial species

The distributions of the *d*_N_/*d*_S_ and *pi*_N_/*pi*_S_ from the resampling process for each species pair are provided in Figure S10. Confidence intervals for mean estimations of *d*_N_/*d*_S_ and *pi*_N_/*pi*_S_ are retrieved using the 2.5% −97.5% quantile from a normal distribution built according to the mean and standard error. The point estimation for species divergence time is estimated from the mean of value (d) from the 500 rounds of resampling. Thus the 95% confidence interval of species divergence time is retrieved using the 2.5% −97.5% quantile from a normal distribution built according to the mean and standard deviation of dS. For the time of transition to sociality, we sampled 10000 sets of observations, each of which consists of a set of values including one observation of (a), (b), (c) and (d) sampled from their distribution built by previous resampling process of 500 random autosomal genes. We then solved the social transition time for each sampled observation set. The point estimation for transition time is obtained from the mean of 10000 sampled calculations. The 95% confidence interval for transition time is retrieved as 2.5%-97.5% quantile from the empirical distribution of 10000 rounds. This will account for all uncertainty of the genome-wide selection strength and species divergence time estimated using the resampling process.

Under the assumption of our calculation for the social period fraction, the social transition time is estimated as the time point when the selection becomes as relaxed as what we observe today between the isolated populations of social species. This level of relaxation in selection intensity would be viewed as the last signal to arise after the completion of social transitions. The social transition times estimated with our *d*_N_/*d*_S_ calculation reflecting sociality have fully evolved based on its impact on purifying selection, despite the start of social transition may be earlier than the estimations.

**Figure M1.**
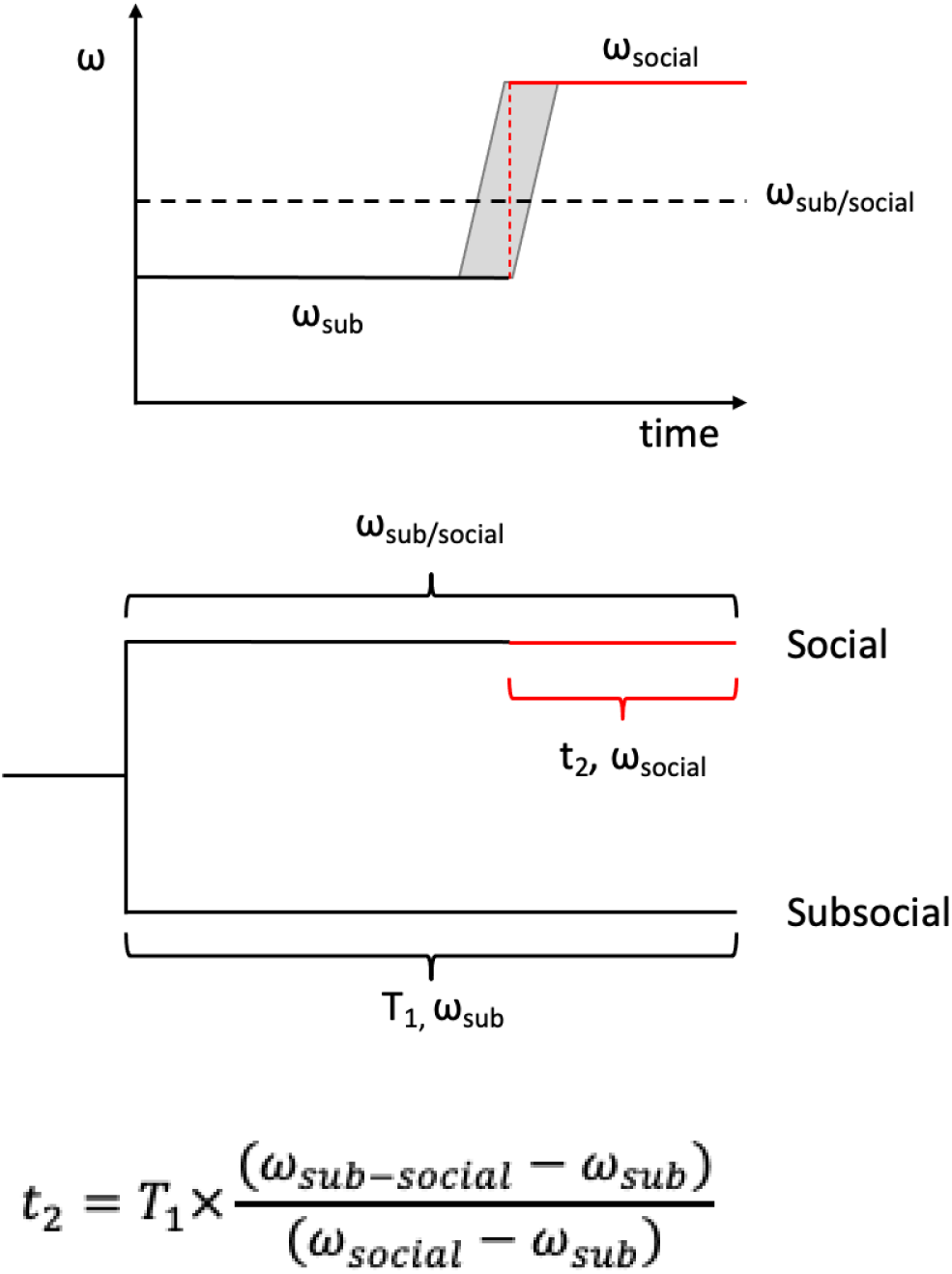
Illustration for calculating the social period length for current social species as an instant process. *t*_2_ denotes the length of the social period. *T*_1_ denotes the divergence time between social species and its sister subsocial species. *ω_sub_* denotes the *d*_N_/*d*_S_ estimated from the subsocial species branch in branch-wise PAML estimations. *ω_sub-social_* denotes the *d*_N_/*d*_S_ estimated from the social species branch in branch-wise PAML estimations. *ω_social_* denotes the *d*_N_/*d*_S_ (In practice, *pi*_N_/*pi*_S_) estimated from the pairwise PAML estimations of two separate social species populations.

### PSMC estimation of social transition time

We used PSMC to reconstruct the historic effective population size for all the six species with a chromosome-level genome. The PSMC method relies on a single diploid individual, where the social species runs out of heterozygosity due to inbreeding. We combine the DNA-seq of two individuals from two separate and diverged populations to retain the loss of heterozygosity.

We benchmarked the fraction of running out of heterozygosity region in the predicted youngest social species *S. dumicola*. We used the GATK pipeline to genotype 9 individuals (5 females and 4 males) from *S. dumicola* and only kept bi-allelic SNP variants among the individuals with a genotype quality filter of 30. We did a sliding window heterozygosity with 100kb window size and 10kb step size on each individual. Windows of an individual without any variants are further classified as run out of homozygosity regions and summarised into ROH fraction by individual (Table S4).

We ran the PSMC method on autosomes and X chromosomes separately. The generation time is fixed to 1 year per generation. The mutation rate used for autosomes is 5e-09 per site per generation and the rate for X chromosomes is scaled to 3.8e-09 with a factor of 0.76 based on the piX/piA being 0.57 instead of 0.75 in subsocial species *S. africanus (Bechsgaard et al. 2019)*. To align the estimated historical Ne estimated between autosomes and X chromosomes, we retrieve all the time points with an estimation of Ne either for autosomal or X chromosomes, the missing value is filled with the linear interpolate according to the closest values on both sides. Then we calculated the historical NeX/NeA as an indicator for evolution of sex-ratio bias. In the end, the curve of NeX/NeA was smoothed using a spline fit in R. The period of accomplished social transition by retrieving the time interval where the decrease of Ne in social species slows down.

### Testing for selection relaxation

Based on the alignment of coding sequences for all filtered autosomal genes, we use HyPhy RELAX to detect the relaxation of intensification of selection on the social species branches. We ran the RELAX test with four different ‘test - reference’ pair settings for the 2303 autosomal genes and 247 X chromosome genes separately. We have one pair that uses all social species terminal branches as the ‘test’ and all subsocial species terminal branches as the ‘reference’ to evaluate the overall effect on selection from the social transition. We have three pairs including each terminal branch of a social species as the ‘test’ and its sister subsocial species terminal branches as well as common ancestor branches as the ‘reference’ to test for relaxed selection in each social transition (Supplementary Figure S3).

## Data access

All of the raw sequencing data was uploaded to NCBI Bioproject PRJNA994315.

The assembled genomes are available in NCBI with the under the following genome accession numbers:

*Stegodyphus dumicola:* CP160598-CP160611
*Stegodyphus tentoriicola:* CP160584-CP160597
*Stegodyphus bicolor:* CP160568-CP160583
*Stegodyphus sarasinorum:* CP160513-CP160528
*Stegodyphus mimosarum:* CP160529-CP160544
*Stegodyphus lineatus:* CP160545-CP160567

Codes for analysis pipeline is available through GitHub: https://github.com/Jilong-Jerome/sociality-in-spiders-dead-end

## Acknowledgement

We thank the support from Independent Research Fund Denmark (Grant number: 0135-00201B). We thank Yael Lubin, Tharina Bird, Virginia Settepani, Lena Grinsted and Clemence Rose for help in collecting samples included in this study.

## Author contribution

Jilong Ma: Analysis design, data analysis, manuscript writing.

Jesper Bechsgaard: Analysis design, manuscript writing.

Anne Aagaard: Analysis design, manuscript refining.

Palle Villesen: Analysis design, manuscript refining.

Trine Bilde: Experiment design, manuscript writing

Mikkel Heide Schierup: Analysis design, manuscript writing, supervision.

