## Supplementary materials and methods for "The genomic consequences and persistence of sociality in spiders"

### Supplementary methods and materials: The genomic consequences and persistence of sociality in spiders

### Section 1 - Sampling and Sequencing

#### DNA - long reads

A single individual from *S. dumicola* (Namibia - Otavi), *S. tentoriicola* (South Africa - Tierberg), *S. mimosarum* (South Africa - Weenen), *S. sarasinorum* (India - Unknown), *S. bicolor* (Namibia - Betta) and *S. lineatus* (Israel - Negev Desert) were sampled, and DNA was extracted using the MagAttract HWM DNA kit from Qiagen (Hilden, Germany). The DNA was fragmented to 15-20 kb fragments using Megaruptor 3, and libraries were prepared using Pacific Biosciences protocol for HiFi library prep using SMRTbell® ExpressTemplate Prep Kit 3.0. Final library was size selected using BluePippin with a 10kb cut-off. Each library was sequenced on three 8M SMRT cells on Sequel II instrument using Sequel II Binding kit 2.2 and Sequencing chemistry v2.0. Loading was performed by adaptive loading, movie time: 30 hours, pre-extension: 2 hours. Between 66 and 99 gb data was obtained (see Supplementary Table S1).

#### Tapestation profiles of DNA extractions for long read sequencing:

Electropherogram of DNA used for PacBio HiFi sequencing on an Agilent TapeStation for each of the six species. The peak around 100bp serves as a standard for both fragment size and intensity.

*Stegodyphus dumicola*:

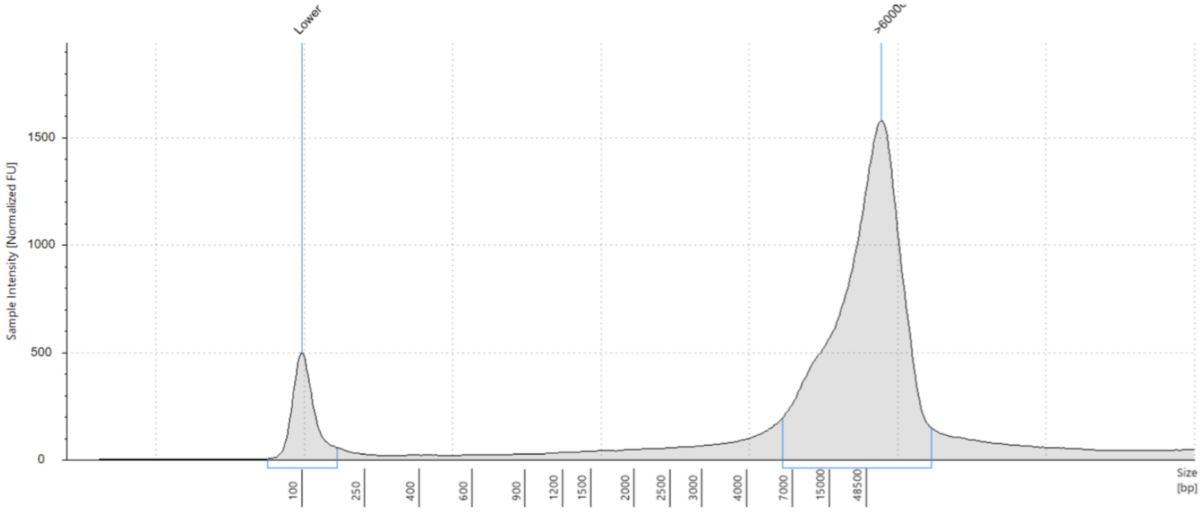


*Stegodyphus tentoriicola*:

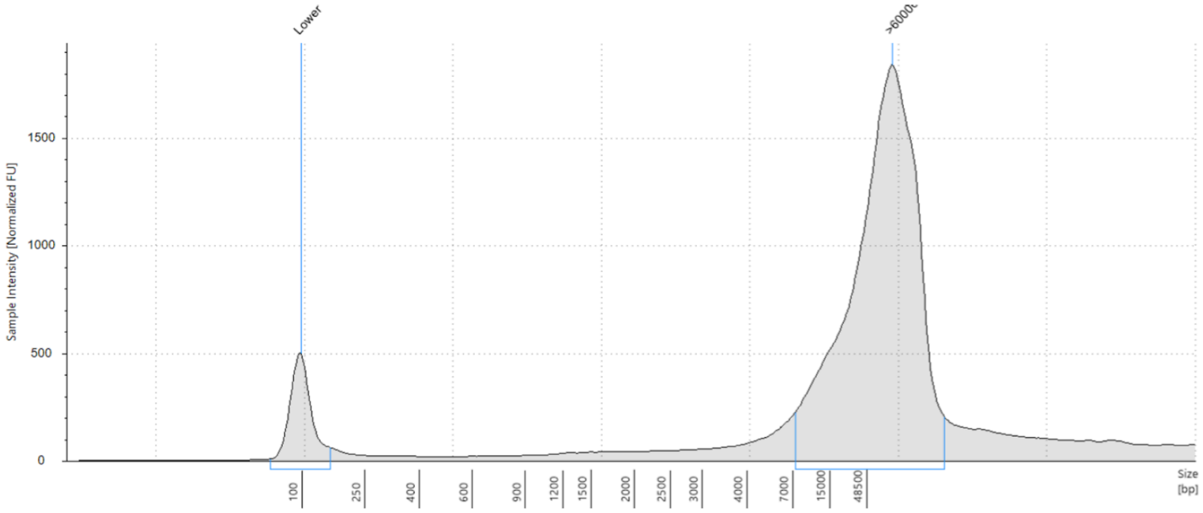


*Stegodyphus mimosarum*:

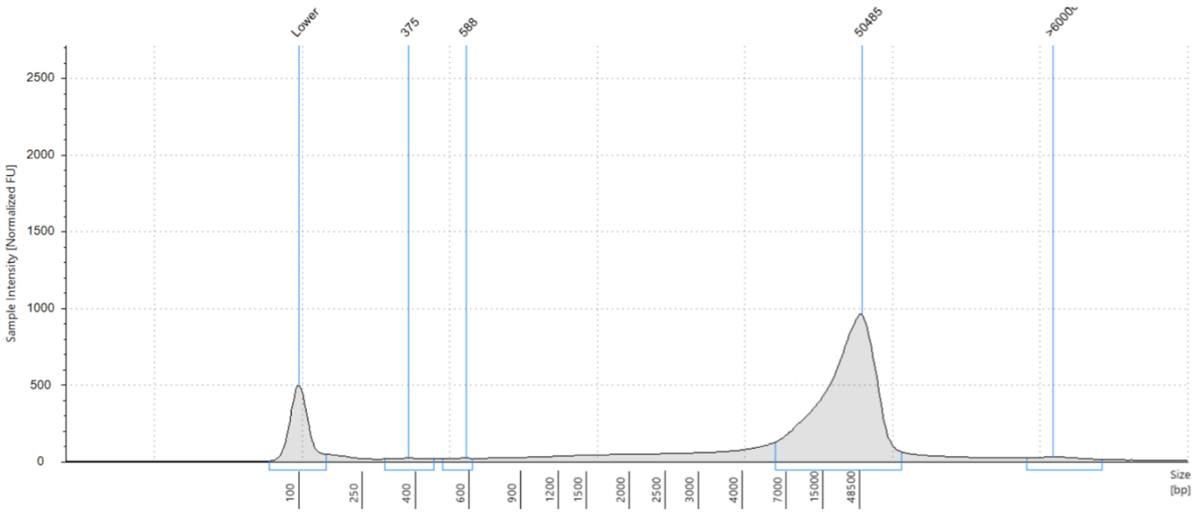


*Stegodyphus sarasinorum*:

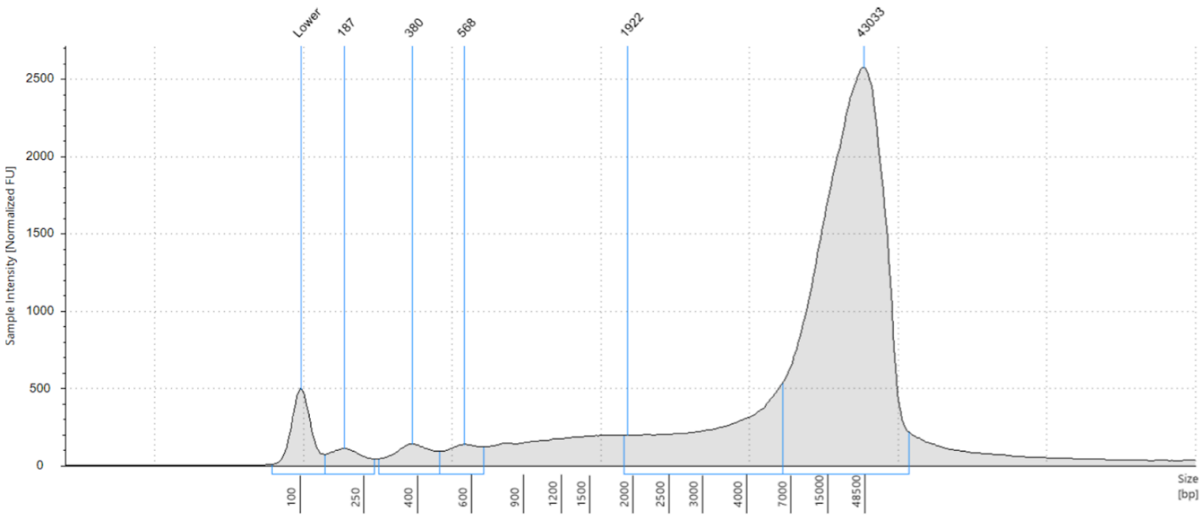


*Stegodyphus bicolor*:

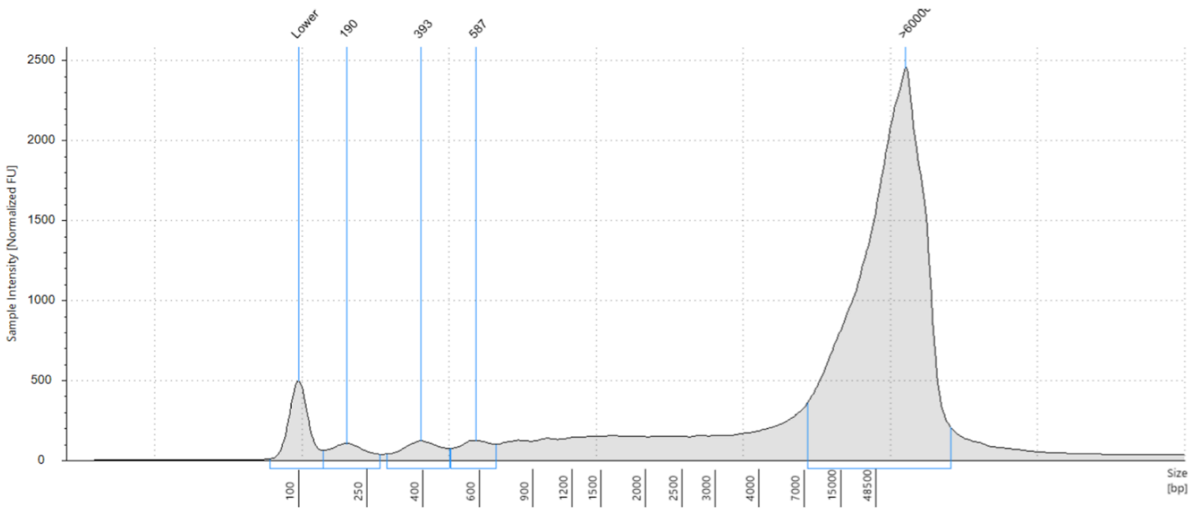


*Stegodyphus lineatus*:

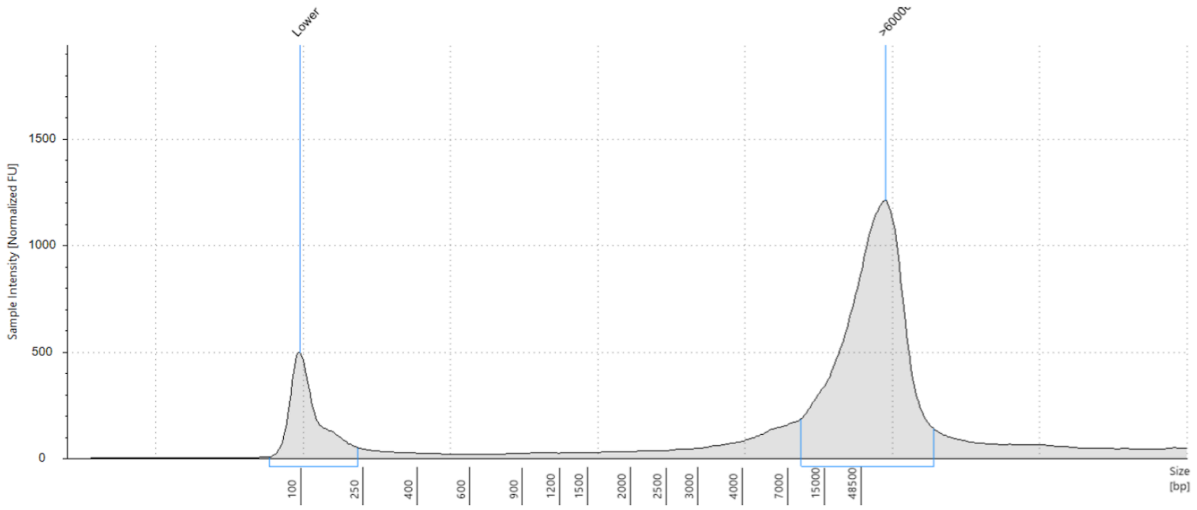


#### DNA - Hi-C

A single individual from each species from the same populations that were sampled for PacBio sequencing were sampled for hi-C sequencing. The social individuals were taken from the same nests. Libraries were prepared from 6 legs per species using the Dovetail® Omni-C® Kit, and each library was sequenced using DNBSEQ-G400 to obtain between 139 and 153 gb 150PE data per sample (see Supplementary Table S1).

### Resequencing

From each of the social species two individuals from isolated genetic lineages were sampled for resequencing to use for estimating ‘social’ dN/dS ratios (piN/piS in practice). From *S. mimosarum*, individuals were sampled in South Africa (Weenen) and Madagascar (Antananarivo); from *S. dumicola*, individuals were sampled from two populations in Namibia (Otavi and Betta); from *S. sarasinorum*, individuals were sampled from Himalaya and Sri Lanka(Settepani, Bechsgaard, and Bilde 2014)(Settepani, Bechsgaard, and Bilde 2014). Five single individuals from the subsocial *S. pacificus* were sampled in India for resequencing to be used for generating a reference genome by aligning reads to chromosome-level assembly of *S. sarasinorum*. DNA from all individuals were extracted using the Qiagen Blood and Tissue kit, Qiagen (Hilden, Germany), and the DNA was sequenced using DNBSEQ-G400 to obtain at between 40 and 146 gb 150PE data per sample (see Supplementary Table S1).

### RNA resequencing

To guide the annotation of protein-coding genes, we sequenced the transcriptomes of several individuals from each species. Three to five families were established by controlled crosses, and three offspring from each family were sampled as spiderlings (n=3), subadults (n=3) and adults  (n=3). All parental individuals came from the same populations as used for long read and Hi-C sequencing. Families were produced as follows: 1) Social species were mated among individuals from the same nest. Initially we created several groups (15+) of 6 subadults to be sure they were virgins until we could determine the sex. When they became adults, we kept only groups with 1 male and 5 females. A few weeks after they were all sexually mature, we froze down the male, and placed the females in individual boxes. The females that laid an egg sac were kept. After the eggs hatched and before matriphagy, one female per group was frozen down and so were six of her offspring. 2) Subsocial individuals were raised from hatching egg sacs produced by females collected in the wild. When sexually mature, females and males with separate mothers were mated (15+), and the male was frozen down. The females that laid an egg sac were kept. After the eggs hatched and before matriphagy, five females were frozen down and so were six of their offspring.

We extracted RNA all individuals using the Qiagen RNeasy Mini kit, and sequencing libraries were constructed using NEBNext Ultra II Directional RNA Library Prep Kit that were sequenced on Illumina NovaSeq 6000 to obtain ~6GB 150PE data per individual. (see Supplementary Table S2)

### Section 2 - *De novo* assemblies and annotations

**De novo assembly**

We generated chromosome-level assemblies for all six species with PacBio HiFi long reads and Hi-C sequencing. We started with using Hifiasm(Cheng et al. 2021)(Cheng et al. 2021), a haplotype-resolved *de novo* assembler for PacBio HiFi reads, to assemble contigs for each species using the default settings. Both PacBio HiFi reads and Hi-C reads were used in this process. We then selected the haplotype with the longer phased assembly graph to retrieve the fasta sequence for contig scaffolding.

Next, we used the Juicer software to align the Hi-C reads to the long read contigs generated in the previous step. We subsequently employed the 3D-DNA (Dudchenko et al. 2017)(Dudchenko et al. 2017) run-asm-pipeline.sh script to order and orient the contigs based on the aligned Hi-C reads. We customize the settings with "-r 0 --editor-repeat-coverage 30 --editor-coarse-stringency 20" to omit mis-join correction rounds in the 3D-DNA scaffolding pipeline. This decision was made because the mis-join correction tends to break long contigs joined from PacBio HiFi long reads in our practice, which would introduce more mis-joining. This customization resulted in a single "mega-scaffold" containing ordered and oriented contigs, with 500 bp Ns introduced at the joints.

Finally, we examined the mega-scaffold Hi-C contact map to identify contigs belonging to the same chromosome, as they exhibited distinct intra- and inter-chromosomal Hi-C contact patterns. We then manually reviewed, edited, and split the mega-scaffold into chromosome-level scaffolds for each species using Juicebox(Durand et al. 2016)(Durand et al. 2016), ultimately generating the final chromosome-level fasta file. The final Hi-C contact maps for all six species are shown in Figure S1. The Hi-C contact map indicates that the scaffolding of chromosomes are visually clean with very few minor misjoined contigs between chromosomes. Further manual curation on the HiC-scaffolding could have been possible to "solve" the few seemingly mis-joints but we should claim clearly that there is no robust way in confirming the precision of the manual curation result. The inherent characteristics of larger contigs derived from long-read HiFi sequencing ensure that the credibility of subsequent analyses focused on local variants and genes remains intact. This is due to the fact that potential local misjoins, typically occurring at a broader spatial scale, do not significantly impact these analyses.

**Genome Annotations**

RepeatModeler2 (Flynn et al. 2020)(Flynn et al. 2020)was initially applied to construct repeat databases that are specific to each species. Following this, we employed RepeatMasker (Tarailo-Graovac and Chen 2009)(Tarailo-Graovac and Chen 2009) to soft-mask the genome assembly for each species, by integrating each species-specific repeat database and the Repbase Arthropoda repeat database(Bao, Kojima, and Kohany 2015)(Bao, Kojima, and Kohany 2015) to form the repeats library.

We used STAR(Dobin et al. 2013)(Dobin et al. 2013) to do spliced alignments for the RNA sequence from every individual sample. The aligned RNA bam files from all the individual samples for a species were then combined using Samtools. This collective data served as transcriptome hints for predicting genes.

We expedited the annotation process by running the BRAKER2(Brůna et al. 2021)(Brůna et al. 2021) ETP mode pipeline independently on each repeat-masked chromosome for each species in parallel. This process involved the use of the aligned species RNA bam file and the NCBI *S. dumicola* protein sequence as the transcriptome evidence and protein homology evidence respectively. The annotations derived from each chromosome were then assembled and merged into a single annotation file in the gff3 format for each species. Lastly, we used BUSCO(Simão et al. 2015)(Simão et al. 2015) to gauge the completeness of the genome annotations, employing the Arthropoda ortholog database for this purpose.

The above annotation pipeline was completed for the majority of the chromosomes in all species, with two exceptions where the BRAKER2 pipeline using ETP mode failed to annotate the local part of the genome. The HiC_scaffold_11 (dum_8) of *S. dumicola* is annotated with the BRAKER2 with only RNA transcriptome data as the evidence. For the ending half of HiC_scaffold_16 (mim_6) of *S. mimosarum*, we used blat to search for *S. bicolor* mRNA sequence against the part of the genome sequence missing annotations. The hits of the blat search were further parsed as the hints for AUGUSTUS gene prediction. The results from AUGUSTUS gene prediction were combined with the results from BRAKER2 ETP mode.

**Sex Chromosome Identification**

We used bwa-mem2 to align reads from a single male individual to the reference genome for each species with a genome assembled. The read depth at each position, covered by at least one read, was obtained using samtools depth. Subsequently, the depth distribution across each scaffolded chromosome was visualized. Chromosomes exhibiting a relative depth in mean and median of half compared to others were identified as X chromosomes, as detailed in Figure S8.

**Genome Quality Control**

To assess genome quality, we employ Mercury, which measures k-mer completeness and base pair accuracy. In the case of subsocial species, where we successfully reconstruct both haplotypes,

Merqury's metrics are provided for the diploid genome assembly. Conversely, for social species characterized by extensive inbreeding, leading to the acquisition of a single haplotype, Merqury's metrics are presented exclusively for the haploid assembly.

### Section 3 - dN/dS ratio estimations

**Benchmarking for substitution of "social dN/dS" with piN/piS within social species**

The general application of dN/dS aims to test the selection strength by investigating the fixed non-synonymous and synonymous substitutions between species. We used piN/piS between divergent populations of the same species as an approximation of "social dN/dS". This implication has a risk of overestimating the dN/dS. As dN/dS compares the fixed nucleotide difference between species under certain selection strength, while piN/piS includes the nucleotide differences that are not fixed by selection yet. Thus, the sites included in the piN/piS would likely contain more deleterious non-synonymous mutations that are not removed by selection, which leads to an overestimated dN/dS. The overestimation could happen if the selection strength on the non-synonymous sites are much higher than the selection strength on synonymous sites, which result in a higher observed piN/piS than the actual dN/dS after fixation.

We benchmarked the reliability of this substitution strategy by comparing the site frequency spectrum of non-synonymous and synonymous sites in an *S. dumicola* population,  which has the shortest population divergent time with potentially more polymorphisms unfixed between populations. We called genotypes for 9 individuals from *S. dumicola* using the GATK pipeline and filtered for bi-allelic single nucleotide variants where the genotype quality is over 30. We used snpEff (version 5.2) (Cingolani et al. 2012)(Cingolani et al. 2012) to build a database with our de novo assemblies and annotations and classify variants as either missense variants or synonymous variants. After the classification, the site frequency spectrum was built for missense variants and synonymous variants respectively (Figure S7).

**Coding gene sequence alignment and filtering for dN/dS estimation**

For getting reliable alignments for dN/dS estimation, we made a strict and conserved filtering for ortholog groups. We ended up analyzing 2302 autosomal genes and 347 single-copy ortholog groups across the 8 species. After retrieving the nucleotide sequence from all species of each ortholog group, we did the alignment using MACSE alignSequences to account for potential frameshift since we are aligning coding sequences(Ranwez et al. 2018)(Ranwez et al. 2018). With MACSE, we ensured that the alignment of each single-copy ortholog is always a multiple of 3 in length.

We do resampling estimation of dN/dS for X chromosomes and autosomes separately. We randomly sample 500 or 100 ortholog groups out of the 2302 autosomal genes or 347 X chromosome genes respectively. The random sample was repeated 500 times for autosomal genes and 100 times for X chromosome genes to get standard error of the mean estimation. For each sampled set of genes, we concatenated the alignment using GoAlign(Lemoine and Gascuel 2021)(Lemoine and Gascuel 2021). We then checked for the concatenated alignment for every codon which contains gaps in any of the species, any codon with gaps in alignment will be marked as removed. The codon marked as removed further divided the whole alignment into continuous alignment blocks in different sizes. An overall distribution of the polymorphic site fraction and alignment block size (Figure S2) suggest that a small size of alignment block with high polymorphic fraction represents local mis-alignment in general. Hence, we filter for alignment blocks that are at least 300 continuous nucleotides long and the fraction of polymorphic sites in a local alignment block should be less than 15%. The above filter was conserved and might lead to fewer sites in final dN/dS estimation, which can be compensated with the concatenation process, but should be more robust to be free from mis-alignment.

**Testing for relaxed selection**

We use RELAX from HyPhy (Wertheim et al. 2015)(Wertheim et al. 2015) to test for the observed higher dN/dS in social species due to intensification of positive selection or relaxed purifying selection. We used the concatenated alignment of all autosomes and X chromosomes separately and 4 different grouping strategies of testing branches and reference branches. For each social species, we used the social species branch as the "Test" and the subsocial sister branches and common ancestor branches as the "Reference". We also have an extra contrasting group where all the social species branches are used as the "Test" and rest branches in the phylogeny are used as the "Reference". The results are shown in Figure S3, we find significant testing results for relaxed purifying selection in autosomes for all contrasting groups.

### Supplementary Figures

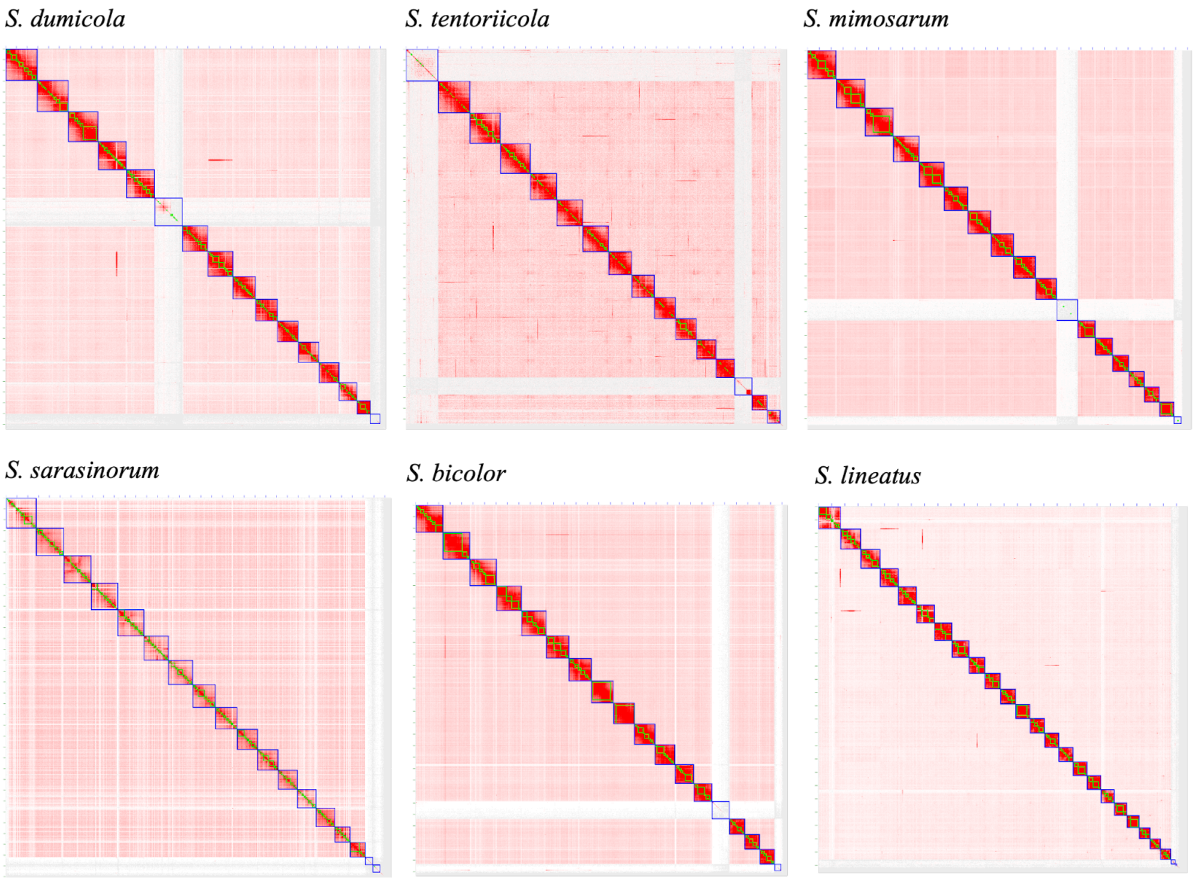


Figure S1. HiC contact map of genome assemblies. The density of red color denotes the hic contact density between regions in the assembly. The green box denotes initial contigs assembled from Pacbio HiFi long reads using hifiasm. The blue box denotes the candidate chromosome-level scaffolds. Each blue box supported by higher intra-scaffold density of HiC contact pattern compared to inter-scaffold HiC contact is identified as a chromosome. The blue box without high intra-scaffold density of HiC contact remains scaffolded.

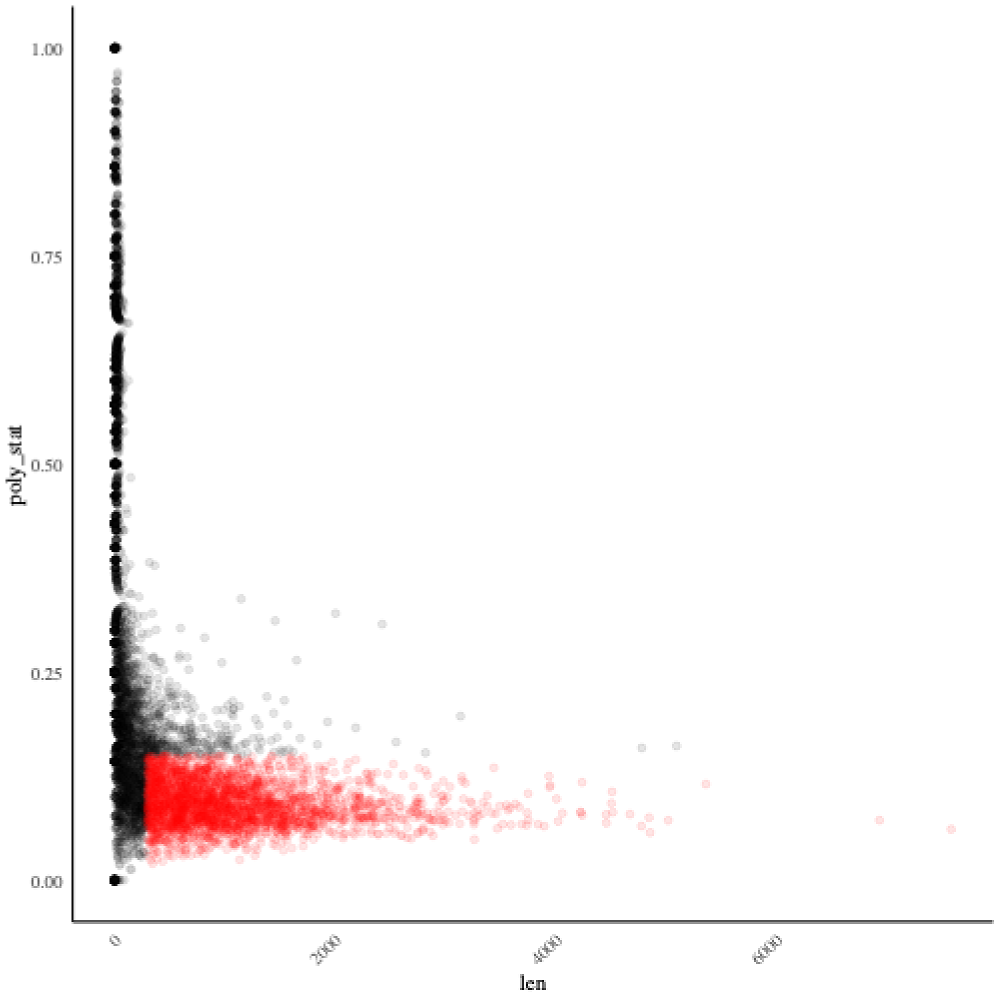


Figure S2 The size of alignment blocks without gaps versus the fraction of polymorphic sites for all alignment blocks of concatenated 2302 autosome gene alignments. The alignment blocks colored in red are further used for estimating dN/dS in PAML(Yang 2007)(Yang 2007).

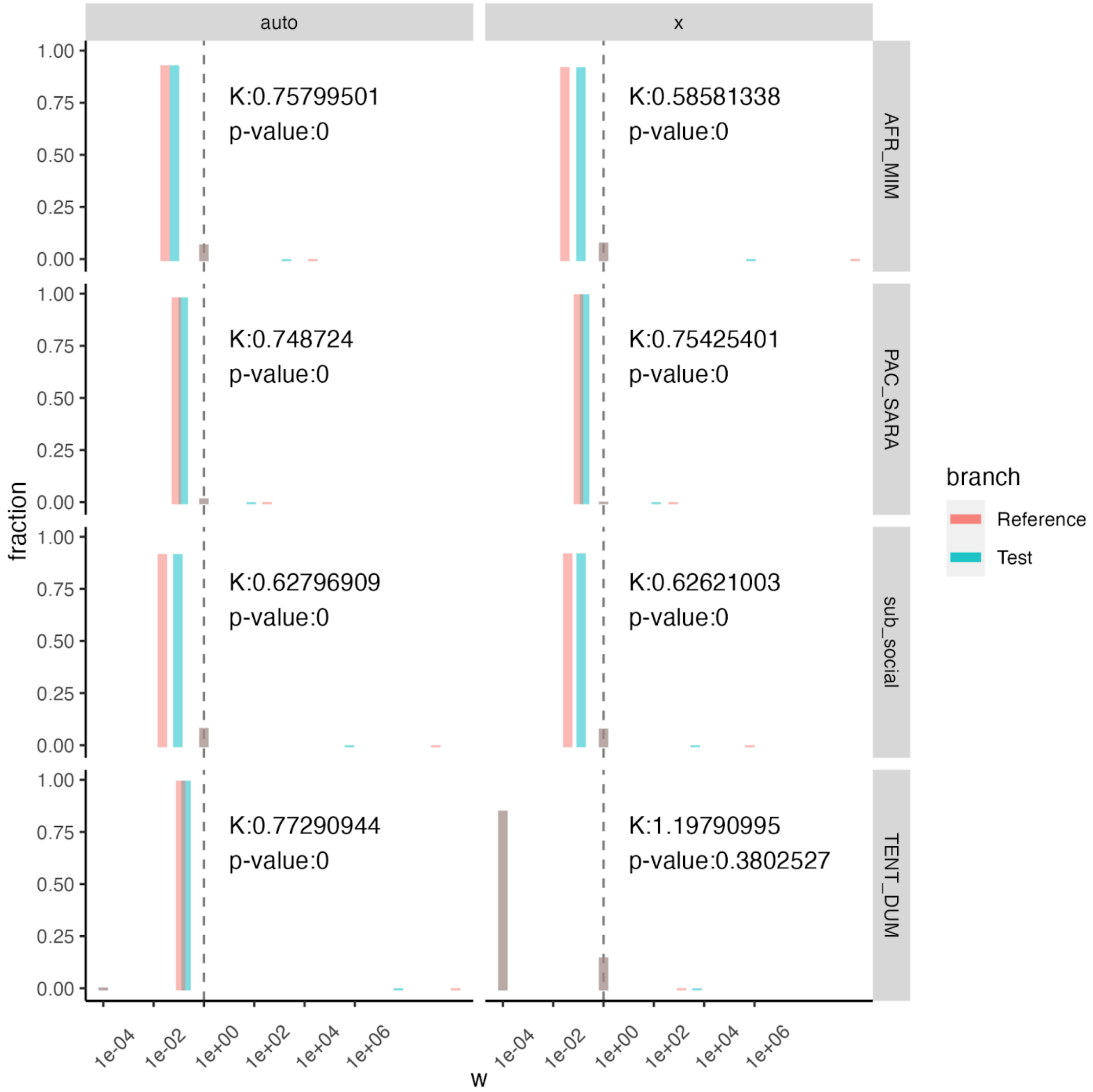


Figure S3. RELAX results for different contrasting pairs in autosomes and X chromosomes separately. K > 1 implies intensification of positive selection and K < 1 implies relaxation of purifying selection. The x axis shows estimated w (dN/dS) and the y axis shows the fraction of sites sharing the corresponding estimated w (dN/dS)

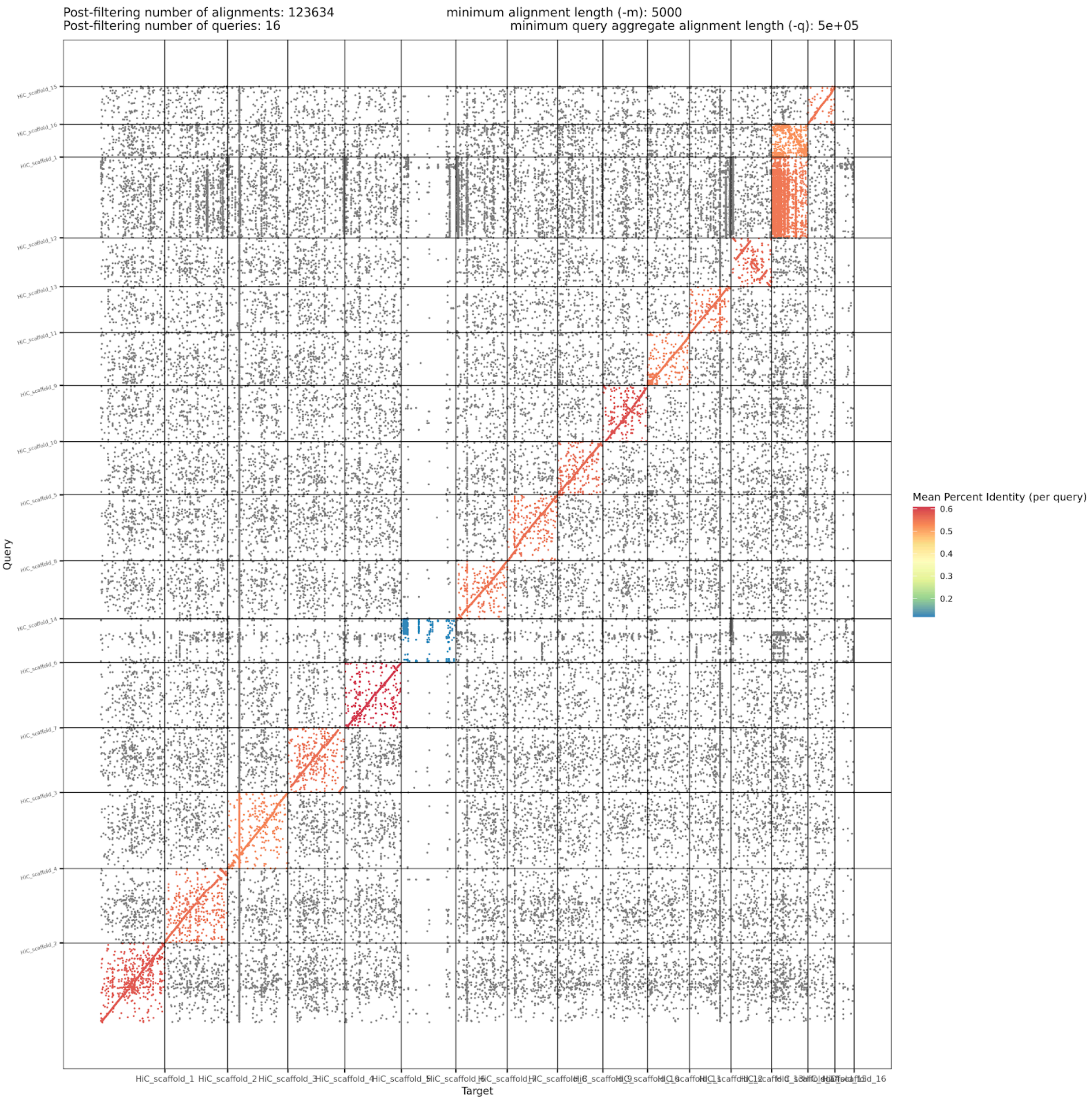


Figure S4. The dot plot from dotPlotly between genome assemblies of *S. dumicola* (x-axis) and *S. tentoriicola* (y-axis). Each point denotes an alignment length of minimum 5000 base pairs. Percentage of identity is shown for each comparison group of chromosomes.

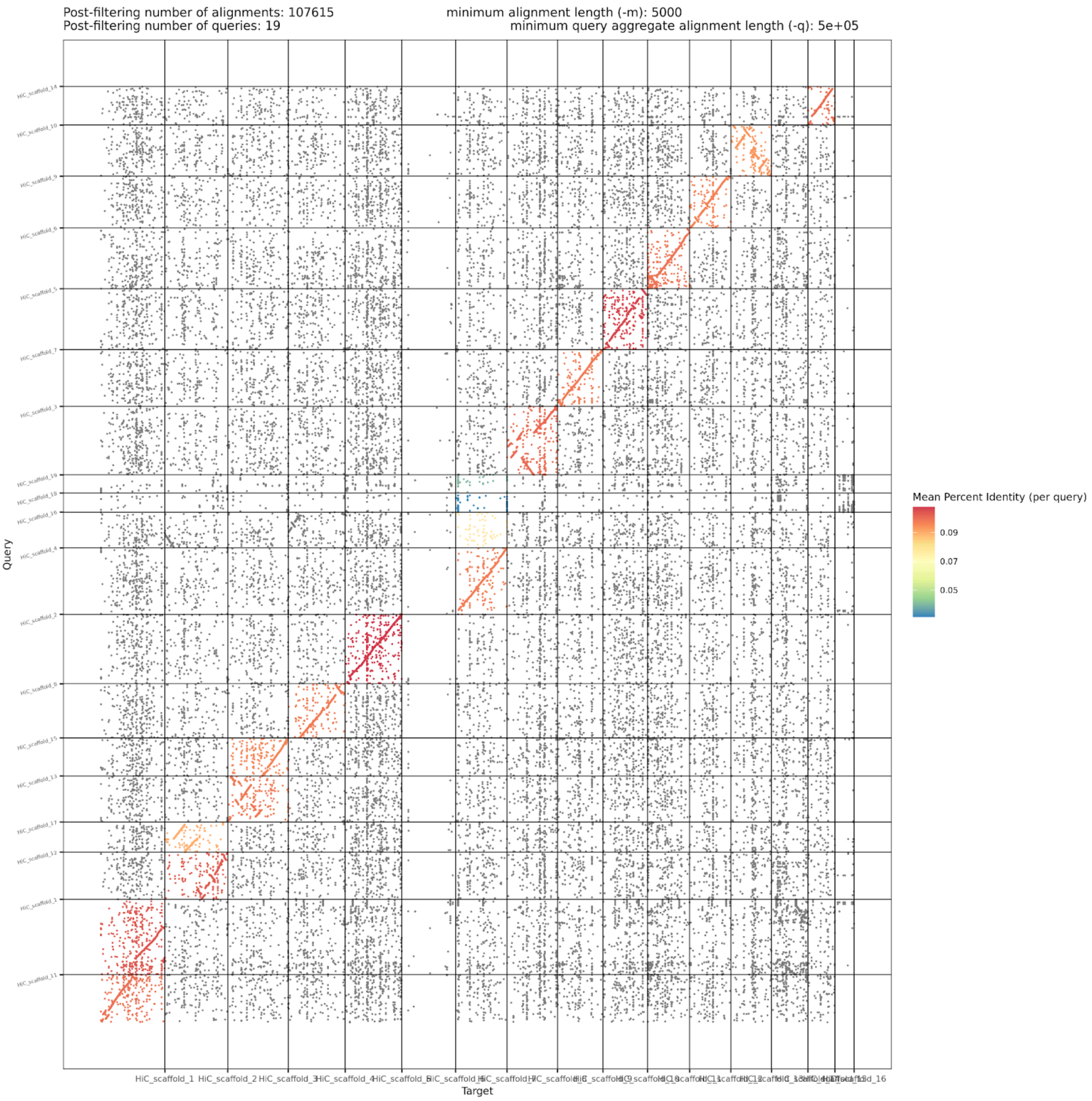


Figure S5. The dot plot from dotPlotly between genome assemblies of *S. dumicola* (x-axis) and *S. sarasinorum* (y-axis). Each point denotes an alignment length of minimum 5000 base pairs. Percentage of identity is shown for each comparison group of chromosomes.

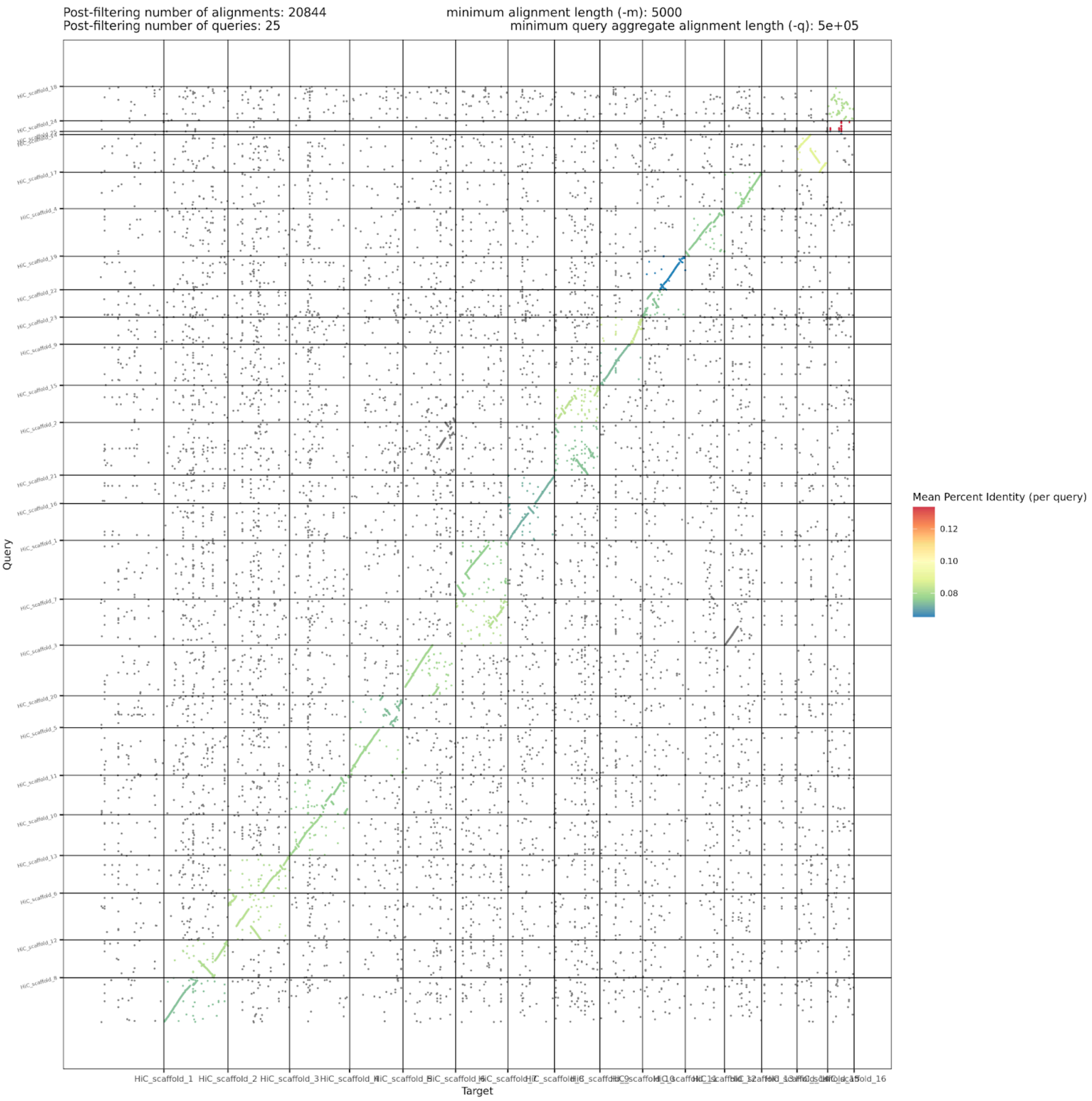


Figure S6. The dot plot from dotPlotly between genome assemblies of *S. tentoriicola* (x-axis) and *S. lineatus* (y-axis). Each point denotes an alignment length of minimum 5000 base pairs. Percentage of identity is shown for each comparison group of chromosomes.

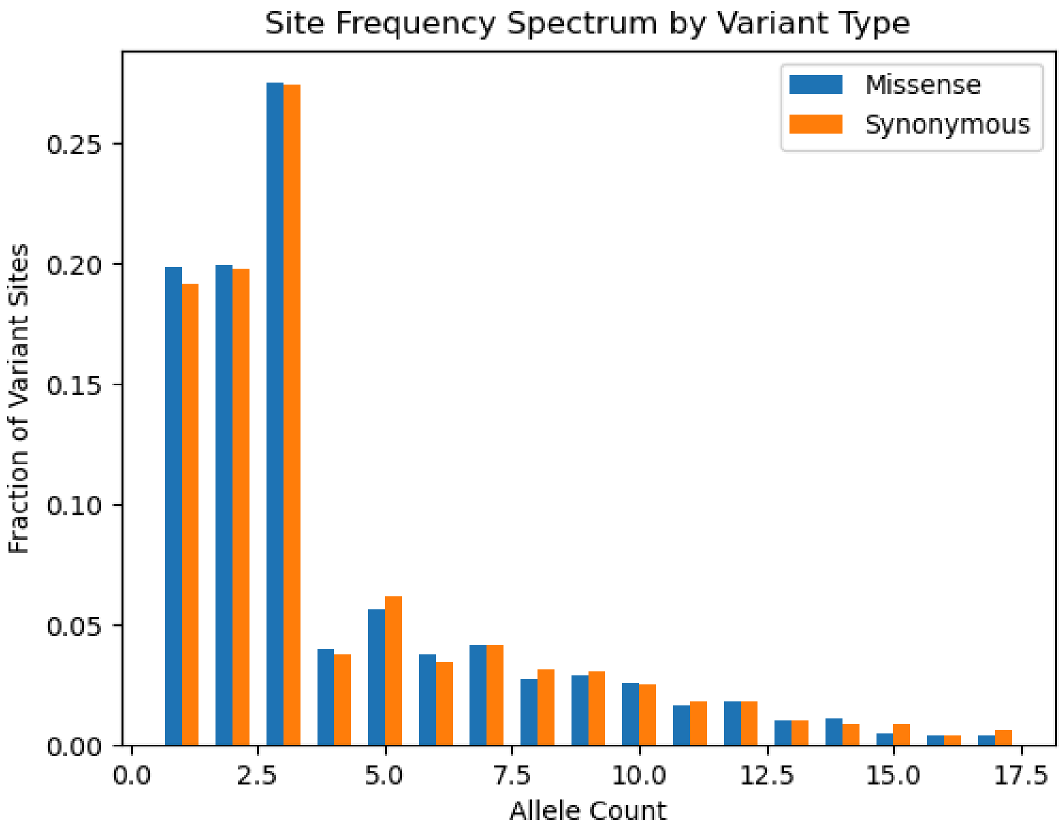


Figure S7. Site frequency spectrums for missense variants and synonymous variants separately based on 9 individuals from *S. dumicola.*

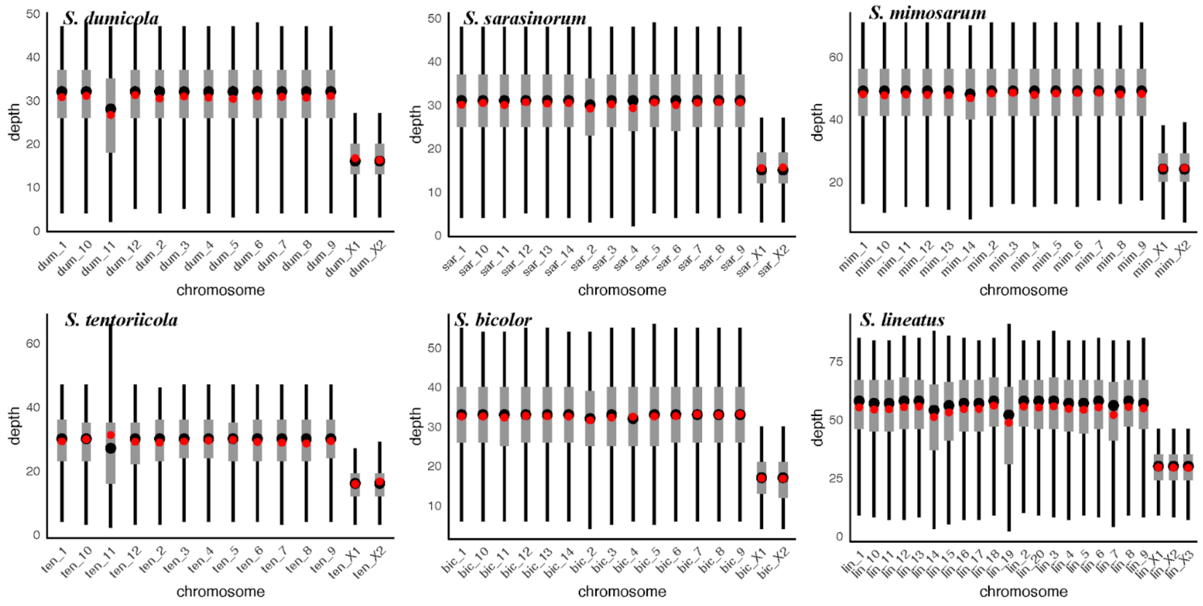


Figure S8. A boxplot of chromosome-level depth distribution from a single male individual for each species. The black lines mark from quantile 2.5% to quantile 97.5%. The grey boxes mark from quantile 25% to quantile 75%. The black points and red points mark the median depth and mean depth respectively.

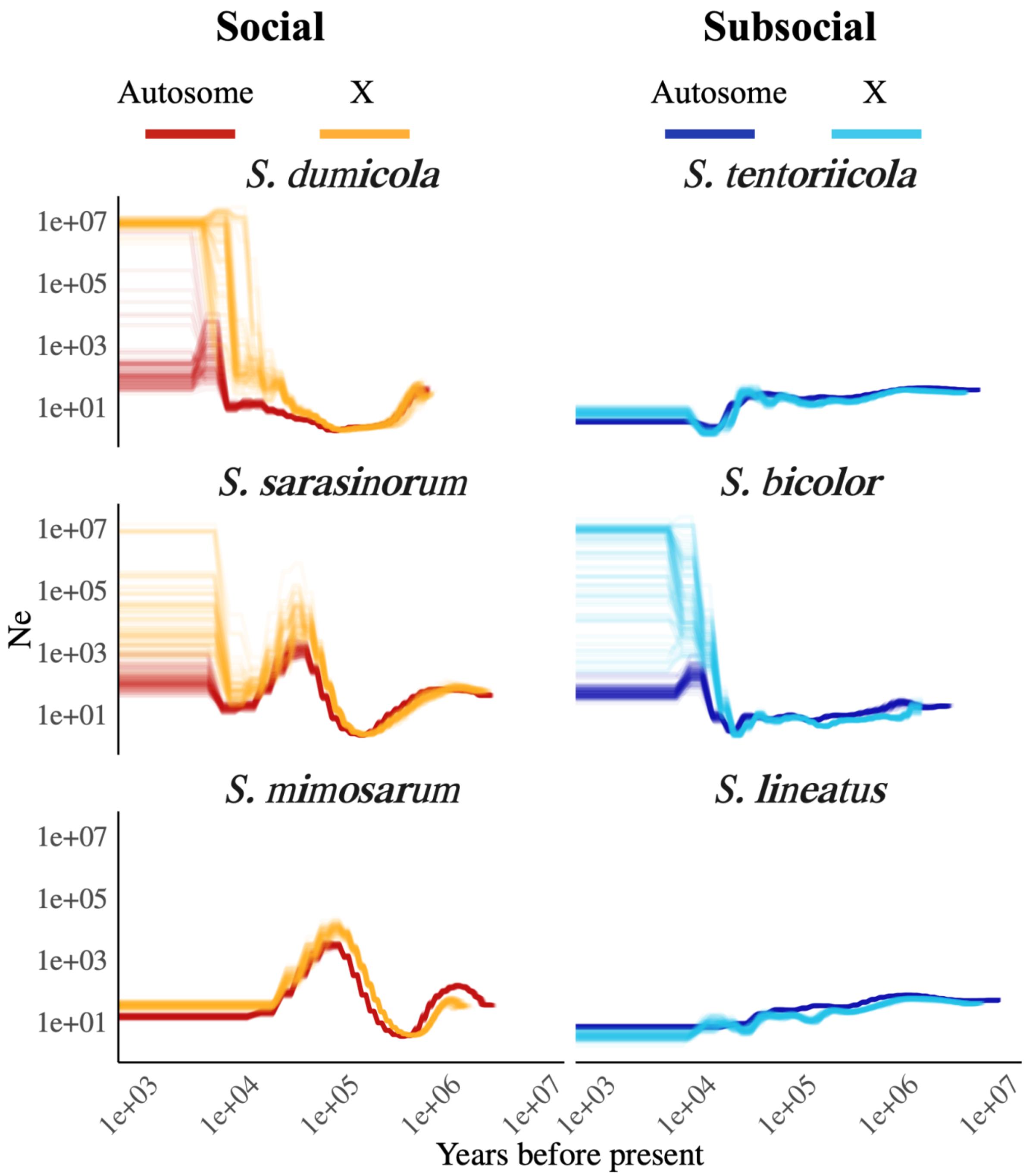


Figure S9. The historical effective population size inferred from the Pairwise Sequentially Markovian Coalescent (PSMC) model with 100 rounds of bootstrapping, setting segment size of 100000bp in resampling process, for different *Stegodyphus* species with chromosome-level assembly. Results from autosomes and X chromosomes are shown separately for each species

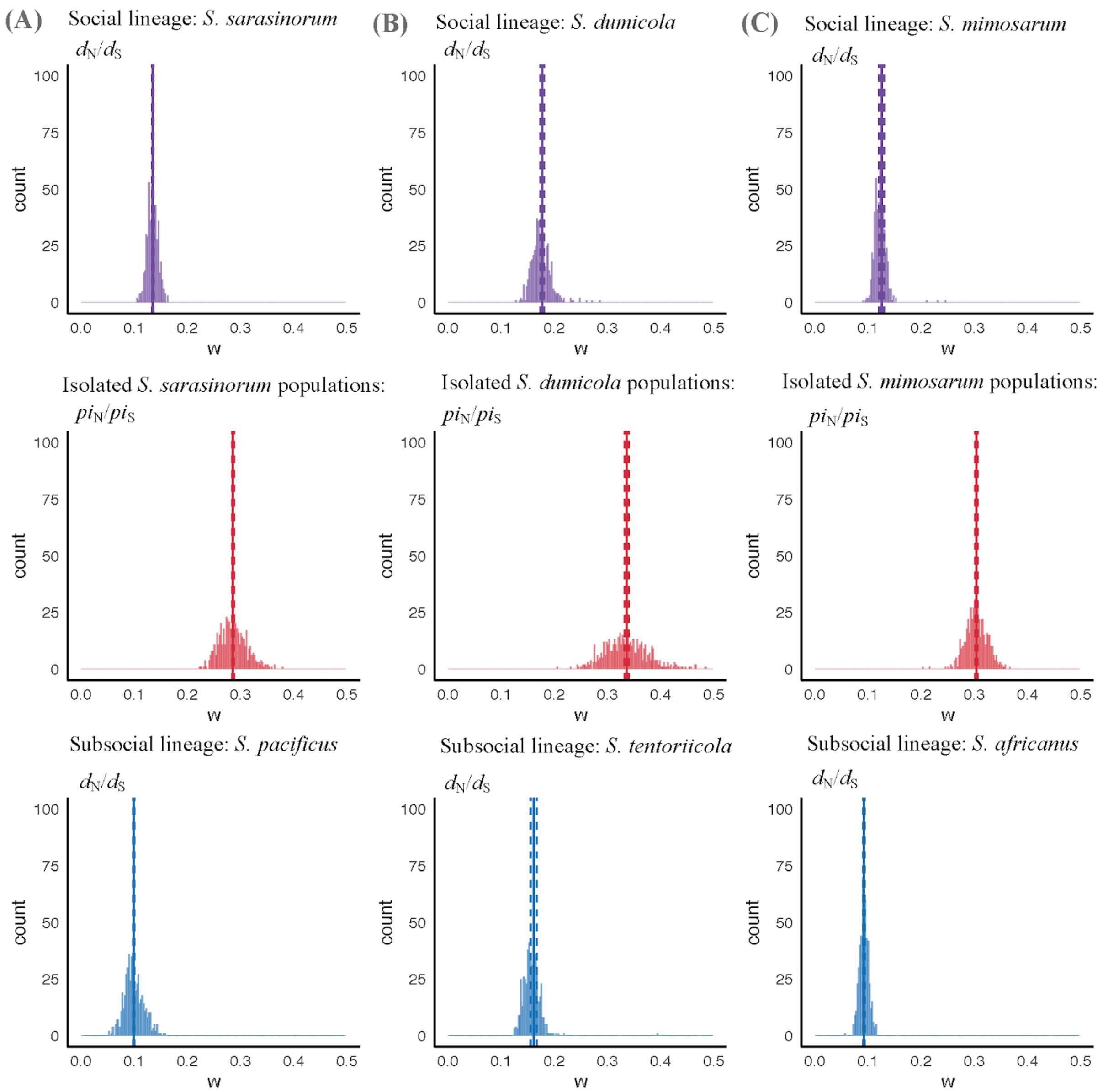


Figure S10. The distributions of dN/dS and piN/piS in each species, based on 500 independent resampling runs, where each run samples random 500 genes out of 2303 autosomal genes without replacement. The point estimation from the mean and 95% confidence interval of the mean are denoted by vertical lines in solid lines and dash lines respectively.

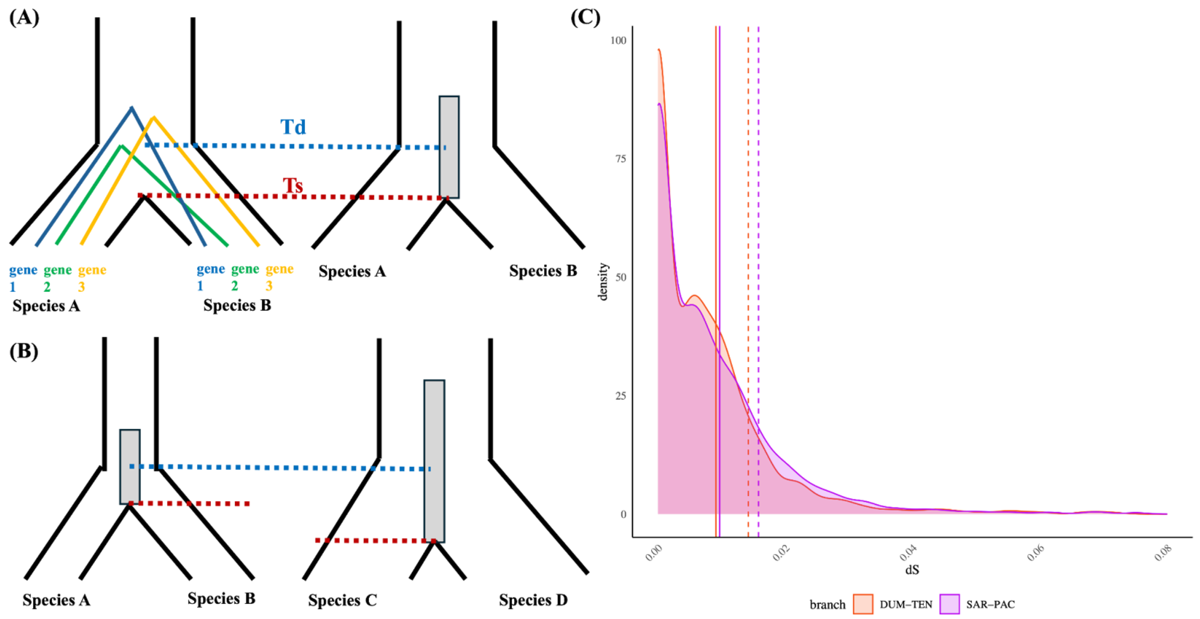


Figure S11. Coalescence expectations of distance between species divergence time (Td) and speciation time (Ts) for *S. dumicola* and *S. sarasinorum*. (A) Different ortholog genes can coalesce at different time points between a pair of species. This provides a time interval on average of 2Ne generations for a gene to coalesce in the ancestral populations, which is the distance between species divergence time (Td) and speciation time (Ts) (from the speciation to the middle point of the grey box). (B) Difference in expected coalescence time between Td and Ts for different ancestral population size. The species pair of species A and species B has a smaller ancestral Ne than the pair of species C and species D. (C) Distribution of dS from each autosomal ortholog gene (2303 in total) for *S. dumicola* (red) and *S. sarasinorum* (pink). The solid vertical lines are the median of the dS distributions, which is expected to correlate with Td (blue dashed line) in (A) and (B). The dashed vertical lines are the 75% quantile of the dS distributions. The distance between the solid vertical line and dashed vertical line in each species is expected to correlate with the width of the distribution, which represents the height of the grey box illustrated in (B).

### Supplementary Tables Text

Table S1. Metadata of sequenced individuals for DNA sequencing (Pacbio HiFi, HiC and short DNAseq)

Table S2. Summary of genome quality for de novo assemblies.

Table S3. Metadata of sequenced individuals for RNA sequencing.

Table S4. Summary of run out of heterozygosity (ROH) regions in *S. dumicola*.

### Supplementary Reference


Bao, Weidong, Kenji K. Kojima, and Oleksiy Kohany. 2015. "Repbase Update, a Database of Repetitive Elements in Eukaryotic Genomes." *Mobile DNA* 6 (June):11.

Brůna, Tomáš, Katharina J. Hoff, Alexandre Lomsadze, Mario Stanke, and Mark Borodovsky. 2021. "BRAKER2: Automatic Eukaryotic Genome Annotation with GeneMark-EP+ and AUGUSTUS Supported by a Protein Database." *NAR Genomics and Bioinformatics* 3 (1): lqaa108.

Cheng, Haoyu, Gregory T. Concepcion, Xiaowen Feng, Haowen Zhang, and Heng Li. 2021. "Haplotype-Resolved de Novo Assembly Using Phased Assembly Graphs with Hifiasm." *Nature Methods* 18 (2): 170–75.

Cingolani, Pablo, Adrian Platts, Le Lily Wang, Melissa Coon, Tung Nguyen, Luan Wang, Susan J. Land, Xiangyi Lu, and Douglas M. Ruden. 2012. "A Program for Annotating and Predicting the Effects of Single Nucleotide Polymorphisms, SnpEff: SNPs in the Genome of Drosophila Melanogaster Strain w1118; Iso-2; Iso-3." *Fly* 6 (2): 80–92.

Dobin, Alexander, Carrie A. Davis, Felix Schlesinger, Jorg Drenkow, Chris Zaleski, Sonali Jha, Philippe Batut, Mark Chaisson, and Thomas R. Gingeras. 2013. "STAR: Ultrafast Universal RNA-Seq Aligner." *Bioinformatics* 29 (1): 15–21.

Dudchenko, Olga, Sanjit S. Batra, Arina D. Omer, Sarah K. Nyquist, Marie Hoeger, Neva C. Durand, Muhammad S. Shamim, et al. 2017. "De Novo Assembly of the Aedes Aegypti Genome Using Hi-C Yields Chromosome-Length Scaffolds." *Science* 356 (6333): 92–95.

Durand, Neva C., James T. Robinson, Muhammad S. Shamim, Ido Machol, Jill P. Mesirov, Eric S. Lander, and Erez Lieberman Aiden. 2016. "Juicebox Provides a Visualization System for Hi-C Contact Maps with Unlimited Zoom." *Cell Systems* 3 (1): 99–101.

Flynn, Jullien M., Robert Hubley, Clément Goubert, Jeb Rosen, Andrew G. Clark, Cédric Feschotte, and Arian F. Smit. 2020. "RepeatModeler2 for Automated Genomic Discovery of Transposable Element Families." *Proceedings of the National Academy of Sciences of the United States of America* 117 (17): 9451–57.

Lemoine, Frédéric, and Olivier Gascuel. 2021. "Gotree/Goalign: Toolkit and Go API to Facilitate the Development of Phylogenetic Workflows." *NAR Genomics and Bioinformatics* 3 (3): lqab075.

Ranwez, Vincent, Emmanuel J. P. Douzery, Cédric Cambon, Nathalie Chantret, and Frédéric Delsuc. 2018. "MACSE v2: Toolkit for the Alignment of Coding Sequences Accounting for Frameshifts and Stop Codons." *Molecular Biology and Evolution* 35 (10): 2582–84.

Settepani, V., J. Bechsgaard, and T. Bilde. 2014. "Low Genetic Diversity and Strong but Shallow Population Differentiation Suggests Genetic Homogenization by Metapopulation Dynamics in a Social Spider." *Journal of Evolutionary Biology* 27 (12): 2850–55.

Simão, Felipe A., Robert M. Waterhouse, Panagiotis Ioannidis, Evgenia V. Kriventseva, and Evgeny M. Zdobnov. 2015. "BUSCO: Assessing Genome Assembly and Annotation Completeness with Single-Copy Orthologs." *Bioinformatics* 31 (19): 3210–12.

Tarailo-Graovac, Maja, and Nansheng Chen. 2009. "Using RepeatMasker to Identify Repetitive Elements in Genomic Sequences." *Current Protocols in Bioinformatics / Editoral Board, Andreas D. Baxevanis ... [et Al.]* Chapter 4 (March):4.10.1–4.10.14.

Wertheim, Joel O., Ben Murrell, Martin D. Smith, Sergei L. Kosakovsky Pond, and Konrad Scheffler. 2015. "RELAX: Detecting Relaxed Selection in a Phylogenetic Framework." *Molecular Biology and Evolution* 32 (3): 820–32.

Yang, Ziheng. 2007. "PAML 4: Phylogenetic Analysis by Maximum Likelihood." *Molecular Biology and Evolution* 24 (8): 1586–91.

Bao, Weidong, Kenji K. Kojima, and Oleksiy Kohany. 2015. "Repbase Update, a Database of Repetitive Elements in Eukaryotic Genomes." Mobile DNA 6 (June):11.

Brůna, Tomáš, Katharina J. Hoff, Alexandre Lomsadze, Mario Stanke, and Mark Borodovsky. 2021. "BRAKER2: Automatic Eukaryotic Genome Annotation with GeneMark-EP+ and AUGUSTUS Supported by a Protein Database." NAR Genomics and Bioinformatics 3 (1): lqaa108.

Cheng, Haoyu, Gregory T. Concepcion, Xiaowen Feng, Haowen Zhang, and Heng Li. 2021. "Haplotype-Resolved de Novo Assembly Using Phased Assembly Graphs with Hifiasm." Nature Methods 18 (2): 170–75.

Cingolani, Pablo, Adrian Platts, Le Lily Wang, Melissa Coon, Tung Nguyen, Luan Wang, Susan J. Land, Xiangyi Lu, and Douglas M. Ruden. 2012. "A Program for Annotating and Predicting the Effects of Single Nucleotide Polymorphisms, SnpEff: SNPs in the Genome of Drosophila Melanogaster Strain w1118; Iso-2; Iso-3." Fly 6 (2): 80–92.

Dobin, Alexander, Carrie A. Davis, Felix Schlesinger, Jorg Drenkow, Chris Zaleski, Sonali Jha, Philippe Batut, Mark Chaisson, and Thomas R. Gingeras. 2013. "STAR: Ultrafast Universal RNA-Seq Aligner." Bioinformatics 29 (1): 15–21.

Dudchenko, Olga, Sanjit S. Batra, Arina D. Omer, Sarah K. Nyquist, Marie Hoeger, Neva C. Durand, Muhammad S. Shamim, et al. 2017. "De Novo Assembly of the Aedes Aegypti Genome Using Hi-C Yields Chromosome-Length Scaffolds." Science 356 (6333): 92–95.

Durand, Neva C., James T. Robinson, Muhammad S. Shamim, Ido Machol, Jill P. Mesirov, Eric S. Lander, and Erez Lieberman Aiden. 2016. "Juicebox Provides a Visualization System for Hi-C Contact Maps with Unlimited Zoom." Cell Systems 3 (1): 99–101.

Flynn, Jullien M., Robert Hubley, Clément Goubert, Jeb Rosen, Andrew G. Clark, Cédric Feschotte, and Arian F. Smit. 2020. "RepeatModeler2 for Automated Genomic Discovery of Transposable Element Families." Proceedings of the National Academy of Sciences of the United States of America 117 (17): 9451–57.

Lemoine, Frédéric, and Olivier Gascuel. 2021. "Gotree/Goalign: Toolkit and Go API to Facilitate the Development of Phylogenetic Workflows." NAR Genomics and Bioinformatics 3 (3): lqab075.

Ranwez, Vincent, Emmanuel J. P. Douzery, Cédric Cambon, Nathalie Chantret, and Frédéric Delsuc. 2018. "MACSE v2: Toolkit for the Alignment of Coding Sequences Accounting for Frameshifts and Stop Codons." Molecular Biology and Evolution 35 (10): 2582–84.

Settepani, V., J. Bechsgaard, and T. Bilde. 2014. "Low Genetic Diversity and Strong but Shallow Population Differentiation Suggests Genetic Homogenization by Metapopulation Dynamics in a Social Spider." Journal of Evolutionary Biology 27 (12): 2850–55.

Simão, Felipe A., Robert M. Waterhouse, Panagiotis Ioannidis, Evgenia V. Kriventseva, and Evgeny M. Zdobnov. 2015. "BUSCO: Assessing Genome Assembly and Annotation Completeness with Single-Copy Orthologs." Bioinformatics 31 (19): 3210–12.

Tarailo-Graovac, Maja, and Nansheng Chen. 2009. "Using RepeatMasker to Identify Repetitive Elements in Genomic Sequences." Current Protocols in Bioinformatics / Editoral Board, Andreas D. Baxevanis ... [et Al.] Chapter 4 (March):4.10.1–4.10.14.

Wertheim, Joel O., Ben Murrell, Martin D. Smith, Sergei L. Kosakovsky Pond, and Konrad Scheffler. 2015. "RELAX: Detecting Relaxed Selection in a Phylogenetic Framework." Molecular Biology and Evolution 32 (3): 820–32.

Yang, Ziheng. 2007. "PAML 4: Phylogenetic Analysis by Maximum Likelihood." Molecular Biology and Evolution 24 (8): 1586–91.
